# Polymer type and surface functionalization determine the impact of nanoplastics on neural crest cell adhesion, migration, proliferation, and apoptosis in primary cultures

**DOI:** 10.64898/2026.11.16.767013

**Authors:** Sylvie Dufour, Redouane Fodil, Frederic Relaix

**Affiliations:** Institut Mondor de Recherches Biomédicales, Faculty of Health, INSERM, Université Paris-Est Créteil, Créteil, France; École nationale vétérinaire d’Alfort, U955 IMRB, F-94700, Maisons-Alfort, France; APHP, Filnemus, Centre de Référence de Pathologie Neuromusculaire Nord-Est-Ile-de-France, Département pathologie, Henri Mondor Hospital, F-94010, Créteil, France

**Keywords:** Nanoplastics, polystyrene, polypropylene, polyethylene, surface functionalization, neural crest cells, cell adhesion, cell migration, cell proliferation, cell apoptosis

## Abstract

Nanoplastics (NP) constitutes a pervasive source of environmental pollution that contaminates all ecosystems, the atmosphere and freshwater. They are absorbed by living organisms and cross biological barriers, with increasing evidence pointing to their harmful effects both at cellular levels and in whole organisms. The impacts of NPs vary not only depending on their size and concentration, but also on the cell type, the polymer composition and surface charge, requiring a comprehensive understanding of the responses of a wide range of cells to these parameters. Because neural crest cells (NCC) are highly proliferative and migratory embryonic progenitors that contribute to a wide range of tissues, they provide a relevant model for investigating the impact of environmental pollutants on developmental cell behaviors. Using a well-established primary NCC culture system, we show that polypropylene (PP) and polyethylene (PE) NPs are more detrimental than polystyrene (PS) NPs to NCC survival and proliferation in a dose-dependent manner. These effects were particularly pronounced for amine-functionalized NPs, which increased NCC apoptosis compared with carboxylate-functionalized NPs. In addition, amine-functionalized PP and PE NPs shortened NCC production period and strongly decrease cell adhesion and migration through alterations of substrate adhesions. Our results indicate that NPs disrupt key biological processes of NCCs, depending on the polymer type and surface functionalization and might contribute to the pathogenesis of structures derived from the NCCs.

## Introduction

In recent decades, human activity has produced large quantities of plastics, including Polystyrene, Polypropylene, polyethylene, Polyethene terephthalate, and Polyvinyl chloride, which offer numerous advantages and have a wide range of applications in the industrial, medical, and household sectors. This has led to the progressive accumulation of plastic waste, representing a major source of environmental pollution and raising increasing concerns about its adverse effects on animal and human health. (1, 2). Throughout their use, recycling, and exposure to environmental stresses, plastics undergo progressive degradation, generating fragments of decreasing size that can reach the nanometer scale (<1 μm). This degradation can also alter their surface properties and increase their capacity to adsorb environmental pollutants. The different chemical and physical characteristics of plastic polymers influence the behavior of micro- and nano-plastics (MNPs) in the environmental and biological systems.

Plastic particles contaminate all aquatic and terrestrial ecosystems, freshwater as well as the atmosphere (3, 4). Ultimately plastics particles enter the body through inhalation, skin contact, and food and liquids ingestion. The small size of MNPs allows them to spread through the bloodstream, cross biological barriers, accumulate and disrupt the organs and tissues functions. (5–8). Over the past decade, a considerable number of studies have sought to analyze the cellular and molecular effects of MNPs on living organisms and cells. MNPs have size-dependent toxicological effects through induction of mitochondrial dysfunction and oxidative stress, generating reactive oxygen species that damage biological macromolecules, altering cellular functions, and leading to inflammation, as well as apoptosis or cellular senescence (7, 9–12). Numerous studies have suggested that MNP exposure may contribute to the development of various pathological conditions, including cardiovascular, neurological, hematopoietic, and urinary disorders, carcinogenesis, dysfunction of the gut–brain axis, and disturbances of the digestive and reproductive systems and fertility (13–18). In addition, it has been shown that MNPs can cross the placental barrier (19, 20) and that maternal exposure to MNPs lead to their accumulation in the placenta and embryonic tissues, with consequences on developmental processes and offspring survival and health (21, 22).

Apart from the whole organism response to exposure to MNPs, which induce specific oxidative stress and inflammation responses, the uptake of particles triggers a large variety of direct cell responses, leading to DNA damage, changes in gene expression and cell fate, endosomal and lysosomal stresses, and alteration of cell polarization, adhesion and migration and survival. These effects vary depending on polymer type, size, concentration, and surface functionalization (23, 24), however, while polystyrene (PS) is not the most prominent plastic pollutant it is the one more often used in cellular studies. Studies performed on macrophages (25), epithelial (26) and endothelial cells (27), as well as on hormone-secreting or migrating neurons (28), and embryonic and neural stem cells (24, 29) have shown that not all cell types respond in the same way to exposure to PS-NPs but less is known about their responses to other types of plastic polymer. Therefore, it is important to characterize the responses of a broader range of cell types to NPs with different polymer type.

Neural crest cells (NCC) are a population of progenitor cells originating from the dorsal neural tube in the early vertebrate embryo and endowed with remarkable stem, proliferative and migratory properties. These features enable the NCCs to colonize various embryonic sites and contribute to a multiplicity of cell types that give rise to a lot of derivatives including cranio-facial structures, melanocytes, cardiac and neuroendocrine derivatives, the neurons and glial cells of the peripheral and enteric nervous system and Schwann cells (30). In addition, NCCs provide large contingents of stem cells in numerous adult tissues. Consequently, disruption of NCC development or function can lead to a broad spectrum of congenital disorders and has also been implicated in various pathological conditions and cancers (31). Their extensive contribution to embryonic development, together with their highly proliferative and migratory properties, makes NCCs a relevant model system for investigating how pollutants, including NPs, may disrupt fundamental cellular processes, such as proliferation, survival, adhesion, and collective migration. In two studies analyzing the effects of in ovo injection of PS-NPs under the vitelline membrane at early stages of chick embryo (32, 33), the authors showed deleterious effects for the embryonic development, associated with heart defects and altered neural tube morphogenesis, as a consequence of the passive targeting of NPs at the dorsal midline of the neural tube, a region where NCCs emerged (32). Cells containing PS-NPs were found protruding into the lumen of the neural tube, a process that was already observed when NCC adhesion was inhibited by peptides known to inhibit integrin-based adhesion (34). However, the impact of NPs on NCC behavior, especially on adhesion, migration, proliferation and survival has not been investigated.

In the present study, we used a short-term primary culture model of embryonic neural tube to investigate the response of NCCs to NPs. We analyzed the effects of 200-nm NPs on NCC adhesion, migration, dispersion, proliferation, and apoptosis, especially the influence of the plastic polymer type, surface functionalization, and doses used on these processes. Our data reveal that NCCs can rapidly internalize PS-NPs and display distinct sensitivity to plastic polymers. Compared to PS-NPs, polypropylene (PP)- and polyethylene (PE)-NPs are more deleterious for NCC dispersion, migration, survival and proliferation, especially for the NPs with amine (NH2) functionalization compared to the carboxylate (COOH) counterparts. In contrast, all types of NPs, even at low doses, affect NCC adhesion, decreasing cell spreading to the substratum, and adhesion sites. Taken together, our results indicate that NPs can profoundly alter NCC behavior in a polymer- and surface modification-dependent manner, with potential consequences for the development and function of structures derived from NCCs.

## Materials and Methods

### Reagents

Bovine plasma fibronectin was from Sigma (cat. no. F1141). Dispase II was from Stemcell Technologies (cat. no. 07913). Amine (NH2)-functionalized nanoplastic (NPs) of polystyrene (PS-NH2), polypropylene (PP-NH2) and polyethylene (PE-NH2) were from Abvigen (cat. no. ABPS-0020-NH2, ABPP-0020-NH2 and ABPE-0020-NH2, respectively). PS-NH2 NPs, coupled to Cy3, were from Molecular Probes (cat. no. F8763). Carboxyl (COOH)-functionalized NPs of polypropylene (PP-COOH) and polyethylene (PE-COOH) were from Abvigen (cat. no. ABPP-0020-COOH and ABPE-0020-COOH, respectively), and polystyrene (PS-COOH), coupled to Cy3, was from Molecular Probes (cat. no. F8810). According to providers technical data, the particle size distribution for F8810 and F8763 were 0.19 ± 0.014 and 0.24 ± 0.010 μm, respectively; for ABPS-0020, ABPP-0020 and ABPE-0020, they were 0.20 ± 0.024, 0.23 ± 0.030, and 0.21± 0.029 μm, respectively. The providers indicate that their plastic particle products are standardized and routine, so they do not measure Zeta Potential, because this parameter is greatly dependent on the medium. The high ionic strength of the culture medium and the presence of fetal bovine serum tend to converge the zeta potential of the carboxylate and amine forms of the NPs toward a narrow and moderately negative value around -10 to -20mV (35). Therefore, a minor increase in the hydrodyamic diameter is expected that we did not quantify.

### Embryos

Quail embryos were used throughout the study. Fertilized eggs purchased from a local farm (La Caille de Chanteloup, Corps Nuds, France) were incubated at 37-38°C until embryos reached the desired developmental stages.

### Primary cell cultures

#### Preparation of neural tube explants for neural crest cell primary cultures

Primary cultures of neural crest cells (NCCs) were produced from trunk neural tubes obtained from quail embryos at different stages covering the NCC delamination and migration process (from embryos with 17 to 25 somite pairs; stage HH13-14 (36)), as described previously (37). After opening the eggshell, the yolk was transferred to PBS. The embryo was cut off from the yolk and an embryo portion of about 750-µm long was excised at the level of the last 5 somites and the nascent pair in the anterior most part of the unsegmented paraxial mesoderm (S1 Fig) with a scalpel under a stereomicroscope and subjected to mild enzymatic digestion using dispase II at 2.5 U/ml for 5 minutes at room temperature. Dispase was washed out and replaced by DMEM, and the neural tube was then separated from the surrounding tissues manually using fine dissection pins under a stereomicroscope, and transferred for 30-60 minutes in DMEM (Sigma, cat. no. D6046) supplemented with 0.5% fetal bovine serum, for recovery from enzyme treatment. Only one neural tube explant per embryo was prepared and was considered as an individual sample.

The neural tubes were explanted onto a variety of substrates depending on the aim of the culture; onto 8-well Lab-Tek® II chambered coverglass (Nunc, cat. no. 155411, hereafter referred to as 8-well plate), in a culture well made from Pyrex cloning cylinder (Corning, cat. no. 11987595) deposited on 12 mm-diameter glass coverslips in 4-well plates (Dutscher, cat. 100040 and 055047, respectively) and hereafter referred to as 4-well plate, or in μ-slide 15-well 3D glass bottom (Ibidi, cat. no. 81507, hereafter referred to as 15-well μ-slide) in 300, 100, or 40 µl of control medium, respectively. Control medium was composed of DMEM containing 100 U/ml penicillin, 100 µg/ml streptomycin, and supplemented with 5% fetal bovine serum. All culture surfaces were pre-coated with fibronectin at a concentration of 10 µg/ml in PBS (approximately 5 µg/cm²) for at least 1 hour at 37 °C. To ensure rapid initiation of NCC migration, the neural tubes were positioned with their dorsal side oriented down toward the substratum. Neural tube explants were cultured at 37°C under normoxic conditions in a 5%-CO2 incubator.

#### Exposure of NCC cultures to nanoplastics

The neural tube explants were incubated for 2 h at 37 °C to allow them to firmly attach to the substratum and for NCCs to begin migrating. Next, depending on the type of the support used, 100, 50 or 10 µl of NP suspensions were gently added to the cultures in 8-well, 4-well plates, or 15-well µ-slides, respectively, and homogenized by gentle pipetting to achieve final volumes of 400, 150 and 50 µl at the desired concentration. For control conditions, the same volumes of control medium were added instead of NP suspensions. Explants were returned to the incubator and analyzed routinely after one day. or after 2 hours in specific experiments as indicated in the figure legends. In one experiment, explants were exposed to 10X PS-NPs for 24 hours, the NP-containing medium was then replaced with control medium, and cultures were maintained for a further 24 hours before fixation. Images of the cultures were acquired on an inverted Nikon Diaphot microscope with 10x and 20x objectives. NP suspensions were prepared extemporaneously in control medium and sonicated for 3 min in a Sonorex TK 30 ultrasonic bath (Bandelin electronic, Berlin, Germany; 50 kHz, 144 W peak ultrasonic power, 36 W effective continuous power) immediately before addition to the cultures, in order to disperse the agglomerates formed during storage (38). Explants were exposed to NP doses spanning three orders of magnitude, from 0.01X to 10X, hereafter referred as 0.01 to 10X concentrations. X corresponds to the number of NPs required to cover the culture area for the various supports used in the study depending on the aim of the experiment and serves as a common factor in the experiments (Table 1 and Eq 1, S1 appendix). This range brackets the internal exposures reported in human tissues, including blood (39) and placenta (40); no equivalent measurement is available for the developing embryo, and the upper doses of the range are therefore supra-physiological by design. The 10X was used to verify the potential short-term toxicity effect of NP accumulation in NCCs.

**Table 1:** Nanoplastics concentrations. Conversion table for 200 nm NP concentrations from X to μg/ml, across the three culture supports used in the study, and for the three NP polymer types, which have different density d (d = 1.03, 0.89 and 0.88 g/ml for PS, PP, and PE, respectively, according to manufacturers). X corresponds to the number of NPs necessary to form one hexagonally close-packed monolayer over the culture area (Eq 1, S1 appendix). Mass concentrations are reported solely to allow comparison with studies expressing doses in µg ml ¹. The final height of medium is the final volume divided by the growth area, and was calculated from Eq 2 (S1 appendix), using densities and final volumes of medium. Because X scales with the culture area, the administered dose per unit area is identical on all three supports, namely 12.5, 10.8 and 10.6 µg cm ² at 1X for PS, PP and PE, respectively. The corresponding mass concentrations differ by up to 1.4-fold between supports because of medium heights. NPs, nanoplastic particles; PS, polystyrene; PP, polypropylene; PE, polyethylene.

| Culture support | Final volume | Growth area | Medium height | X |  |  | Polymer | X |  |  |  |
|---|---|---|---|---|---|---|---|---|---|---|---|
|  |  |  |  | 10<sup>6</sup>NP | 10<sup>6</sup>NP<br>cm<sup>-2</sup> | 10<sup>6</sup>NP<br>ml<sup>-1</sup> |  | 0.01 | 0.1 | 1 | 10 |
|  | µl | cm<sup>2</sup> | mm |  |  |  |  | µg ml<sup>-1</sup> |  |  |  |
| 8-well | 400 | 0.70 | 5.71 | 2020 | 2886 | 5050 | PS | 0.22 | 2.2 | 21.8 | 218 |
|  |  |  |  |  |  |  | PP | 0.19 | 1.9 | 18.8 | 188 |
|  |  |  |  |  |  |  | PE | 0.19 | 1.9 | 18.6 | 186 |
| 4-well | 150 | 0.28 | 5.31 | 816 | 2888 | 5440 | PS | 0.23 | 2.3 | 23.5 | 235 |
|  |  |  |  |  |  |  | PP | 0.20 | 2.0 | 20.3 | 203 |
|  |  |  |  |  |  |  | PE | 0.20 | 2.0 | 20.1 | 201 |
| 15-well<br>µslide | 50 | 0.125 | 4.00 | 360 | 2880 | 7200 | PS | 0.31 | 3.1 | 31.1 | 311 |
|  |  |  |  |  |  |  | PP | 0.27 | 2.7 | 26.8 | 268 |
|  |  |  |  |  |  |  | PE | 0.27 | 2.7 | 26.5 | 265 |

### Cell Proliferation assays

Cell proliferation in culture was monitored using the click-It EdU Cell proliferation kit for imaging from Life Technologies (cat. no. C10638, C10337). Briefly, NCC primary cultures were generated for 19 hours in μ-slide 15-well and EdU was added to the culture medium at the final 20 µM concentration for 1 hour. Immediately after EdU incorporation, medium was removed and cultures were fixed in 4% PFA in PBS for 15 minutes at room temperature, permeabilized in 0.5% Triton-X100 for 15 minutes and treated for EdU detection using Alexa-555 or -488 Fluor azide in accordance with the manufacturer’s guidelines. After DNA staining with Hoechst, cultures were observed using an AxioVert A1 microscope using a LD A-Plan 10x/0.25 or 20x/0.35 Ph1 objectives.

### Immunofluorescence labeling of cultures

The following primary antibodies were used: mouse mAb to paxillin (Clone 165, BD-Transduction Laboratories, 1/100), mouse mAb to Sox-10 (Clone A2, Santa Cruz, 1/200), rabbit Ab to Sox-10 (Bio SB, cat. no. EP268, 1/250), rabbit Ab to anti-activated Caspase3 (Millipore, cat. no. AB3623, 1/100). Filamentous actin was detected using Alexa-Fluor 488 Phalloidin (Molecular probe, cat. no. A12379). NCC primary cultures were performed on the three types of supports. Cultures were fixed in 4% paraformaldehyde (PFA) in PBS for 15 minutes at room temperature for detection of antigens. After permeabilization with 0.5% Triton X-100 in PBS for 5 minutes, cultures were rinsed twice, blocked in PBS-3% BSA and subjected to immunofluorescence labeling using primary antibodies followed by incubation with appropriate secondary Ab conjugated to Alexa-fluor 488, Cy-3 or Cy-5 (Jackson Immunoresearch Laboratories), and processed for DAPI staining to visualize cells’ nuclei before mounting in ImmuMount medium (Shandon). In some experiments Alexa-fluor 488 conjugated WGA (1/200e, cat. no. W11261, Life Technologies) was added to the secondary antibody mix to label cell membranes.

Images were acquired with a Zeiss AxioImager M2 epifluorescence microscope equipped with 10X-63X fluorescence objectives (Acroplan 10X/0.25, 20X/0.45, Plan-Neofluar 40X/0.75 and 63X/1.25 oil) or with a Zeiss LSM 900 confocal microscope equipped with Plan-Apochrome 40X/1.30 oil objective. In some experiments, Z-series were acquired with step size 0.25μm. Data were collected using the Zen or Airyscan2 software and processed using ImageJ software.

### Time-lapse video microscopy

For the analysis of NCC migration front progression, NCC primary cultures were performed in 8-well Lab-Tek® II. Up to 4 neural tubes were distributed into each well, far enough apart for their outgrowths to be imaged separately, and cultured in control medium. Neural tubes were cultured at 37°C in a humidified incubator for about 2 hours until they adhered firmly to the dish and NCCs initiated migration. The NPs were then added to the wells and cultures were placed on the motorized stage of a Leica DMIRE2 microscope equipped with a CoolSNAP HQ camera (Roper Scientific). Time-lapse video microscopy was performed with a 10X objective and phase contrast images were captured every 5 minutes for 18 hours using the Micromanager software. Recording started 60 to 90 minutes after the addition of NPs; this time point is defined as t0.

For the cell tracking, NCC primary cultures were performed in CO2 Independent medium from Gibco (cat. no. 18045-088) containing 100 U/ml penicillin, 100 µg/ml streptomycin, supplemented with 5% serum, 2 mM glutamine, and 1.4 μl/ml of Incucyte® Nuclight Rapid NIR Dye (Essen BioScience, cat. no. 4717) for live cell nuclear labeling. This medium was defined as the control medium in these experiments. Neural tubes were cultured at 37°C for about 2 hours. The NPs were then added to the wells and cultures were placed on the motorized stage of a LSM 900 Airyscan 2 Zeiss confocal microscope into a heated chamber with a humid atmosphere. Recording started 60 to 90 minutes after the addition of NPs; this time point is defined as t0. Nuclei were imaged in parallel for all explants at multiple stage positions, using a 20x objective, every 3 minutes for 18 hours, with ZEN software (version 3.0; Carl Zeiss Microscopy). The nuclear dye (excitation and emission maxima, 650 and 673 nm) was excited with the 640 nm laser line, and its emission was collected between 632 and 700 nm. Brightfield images were captured simultaneously with fluorescence data using T-PMT.

### Quantifications

#### Outgrowth area

For each explant, overlapping phase-contrast images covering the whole NCC outgrowth were acquired at 24 hours with a 10x objective. They were assembled manually to reconstitute in a single image the entire NCC outgrowth using Affinity Photo 2 (version 2.6.5; Serif, Nottingham, UK). On this reconstructed image, the outline of the outgrowth and the length of the neural tube were traced manually with the freehand and straight-line selection tools of ImageJ. To account for differences in neural tube length among the explants and allow comparison between experiments, the measured outgrowth area was multiplied by the ratio of a reference length to the length of the neural tube. The reference length was the mean length of neural tubes measured at 24 hours of culture. Normalized outgrowth areas are expressed in µm². One value was obtained per explant; the explant being the statistical unit for this measurement.

#### Cell spreading and density in the outgrowth

Phase-contrast images of neural tube explant cultures for 24 hours were acquired with a 20x objective at different positions along the long axis of each explant. Images were obtained from at least three explants per condition in at least two independent experiments. Cell spreading was quantified on these images by tracing manually, with the freehand tool of ImageJ, the outline of individual NCCs per image, and measuring their area in µm². Cell density was quantified by counting, with the Cell Counter tool of ImageJ. The NCCs present in square regions of 100 µm side per image and expressed as cells per 10 µm².

#### NCC front progression, migratory parameters and production period

Front progression, individual cell migration, and the duration of NCC production were measured on the video recordings described above (see Time-lapse video microscopy). The position of the migratory front of the NCC population was determined at 1-hour intervals with a custom graphical user interface written in MATLAB (MathWorks, Natick, MA, USA), as previously described (41). Front progression was expressed as the distance covered by the front since t0. For each video, trajectories and positions of nuclei of 10 to 20 NCCs, located at the periphery of the outgrowth on each side of the neural tube, were tracked over 12 hours using MetaMorph (version 7; Molecular Devices, San Jose, CA, USA). For each cell, mean speed was calculated as the total length of the trajectory divided by the tracking duration. Persistence was calculated as the ratio of the straight-line distance between the initial and final positions to the total length of the trajectory. Hourly speed was calculated as the length of trajectory covered during each successive 1-hour interval. NCC production and the onset of their migration were followed along the border of the neural tube on the same recordings, using the transmitted light channel. The duration of NCC production was defined as the time from t0 to the last time point at which NCCs were seen separating from the neural tube on recording. The morphologies of the NCCs were observed throughout video recordings. The moment when any changes in the morphology of the NCCs occurred during migration, relative to time t0, was determined using video recordings. When no significant change in NCC morphology was observed during the 18-h recording the value was set as 18 for each cell analyzed.

#### Cell Proliferation

Fluorescent images of EdU and DAPI stainings were taken with the 20x objective in various zones of NCC outgrowth along the long axis of each explant and the DAPI+/EdU- and EdU+ NCC were counted manually using cell counter tool of ImageJ.

#### Plastic particles incorporation

Confocal z-series were acquired with a Plan-Apochromat 40×/1.40 oil objective and a z-step of 0.25 µm. For each image, 5 to 10 individual cells were selected at random, their outline traced manually on the phase-contrast channel using ImageJ, and the corresponding region cropped from the full z-series. Internalization was quantified for PS-NPs only, since PP- and PE-NPs were not fluorescently labeled. NPs were then counted manually plane by plane along the z-stack. Only NPs located between the basal and the apical optical sections of the traced cell were counted, so that NPs lying on the substrate outside the cell footprint were excluded; NPs possibly located on the plasma membrane could not be distinguished from fully internalized ones and are therefore included in the counts. When NPs formed aggregates, the number of NPs in the aggregate was estimated from its apparent size and from the extent of the fluorescent signal along the three axes, using isolated single NPs imaged in the same acquisition as a reference for the signal of one NP. Counts were performed by a single observer, and images were not blinded to the experimental condition. The number of NCCs analyzed, explants, and independent experiments per condition are indicated in the legend of the corresponding figures.

#### Cell substrate adhesions

Substrate adhesions (SA) were quantified on neural tube explants cultures fixed at 24 hours in 8-well or 4-well plates and immunostained for paxillin and with nuclei labelled with DAPI. For each explant, one to three images were acquired at the periphery of the outgrowth in randomly selected fields, with a Zeiss LSM 900 confocal microscope mounted on an Axio Observer Z1 stand and equipped with a Plan-Neofluar 40×/1.3 oil objective, from two to four explants per condition. Images were analyzed with Fiji (ImageJ2 version 2.16). Each image was segmented by manual thresholding, and individual SAs were detected with the Analyze Particles tool, using a minimum size of 0.2 µm². The area and circularity (4π × area / perimeter²) of each SA were measured, and the number of NCCs in each image was determined by counting DAPI-stained nuclei. For each image, the SA number per NCC was calculated as the number of SAs divided by the number of NCCs, and the total SA area per NCC as the summed area of all SAs of a minimum size of 0.2 µm² divided by the number of NCCs. SA size and circularity were averaged over all SAs of the image. The image was the unit of observation for these measurements.

### Statistical Methods

Statistical analyses were performed with GraphPad Prism (version 10.1.3). Each treated condition was compared with the control condition without NPs, and for some experiments also between the two types of functionalized NPs. Each experimental series included control cultures run in parallel. In some experiments, the culture conditions with NH2- and COOH-NPs shared the same controls. For migration experiments, the control cultures of independent experiments were pooled; each treated condition was then compared with the pooled control. Each NCC outgrowth produced in culture was considered as an individual sample. The unit of analysis differed between measurements and is indicated in each figure legend: the explant for outgrowth area and duration of NCC production; the image for NCC density, EdU, activated caspase-3, and paxillin stainings, and the individual NCCs for NP counts, cell spreading, speed (average and hourly) and persistence.

For comparisons involving more than two groups, normality was assessed in each group with the Shapiro-Wilk test and homogeneity of variances with the Brown-Forsythe test. When all groups met both assumptions, data were analyzed by one-way ANOVA followed by Fisher’s least significant difference (LSD) test for comparisons of each condition with the control. Otherwise, the Kruskal-Wallis test was used, followed by Dunn’s test for comparisons of each condition with the control. In both cases, no correction for multiple comparisons was applied. Comparisons between two groups, namely for front progression and hourly speed at each time point, were performed with unpaired two-tailed t-tests, without correction for multiple comparisons. Differences were considered significant at p < 0.05. Data are presented with mean ± SD or mean ± s.e.m., and indicated in each figure legend. For cell spreading, cell speed, and persistence, data are presented as SuperPlots (42). In these SuperPlots, values for individual NCCs are shown as transparent dots and the mean of each explant as a solid black dot, with bars indicating the mean ± SD calculated on all cells. Explant means are shown for visualization only; statistical tests were performed on all cell values. The numbers of explants, images or cells analyzed, as well as the number of independent experiments are given in each figure legend.

### Disclose use of AI tools

During the preparation of this manuscript, the authors used Claude (Anthropic; model identifier: claude-sonnet-5-5; web interface at claude.ai) to: (i) verify internal consistency across the main text, figure legends, and tables; and (ii) suggest stylistic rephrasing in the Methods, Results, Figure Legends, and Discussion sections to improve clarity and readability. All experimental data were acquired, and all statistical analyses were conducted directly by the authors using GraphPad Prism. No raw or processed data, figures, or images were generated by artificial intelligence. The authors reviewed, validated, and take full scientific responsibility for all content presented in the manuscript.

## Results

### NCCs rapidly incorporate and accumulate Polystyrene nanoplastics

To examine the capacity NCCs to incorporate and accumulate nanoplastics (NPs) we cultured neural tube collected from trunk region of quail embryos at stages of NCC delamination and migration process (S1 Fig). Neural tubes were put in culture in control medium for two hours, then different amounts of polystyrene (PS)-NPs, 0.01, 0.1, 1 and 10X (see Table1), were added. We observed that after two hours of incubation in the presence of NPs, NCCs took up PS-COOH (Fig 1A) and -NH2 NPs (S2A Fig) into their cytoplasm in a dose-dependent manner, the particles being found within the cell outline on confocal z-series. We observed that after two hours of incubation in the presence of NPs, NCCs took up PS-COOH (Fig 1A) and PS-NH2 NPs (S2A Fig) in a dose-dependent manner, the particles being found within the cell outline on confocal z-series. The number of NPs per cell was broadly distributed across the NCC population, reaching up to 5 PS-COOH and 2 PS-NH2 NPs at 0.01X concentration, up to approximately 30 and 9 at 0.1X, and up to approximately 80 and 30 at 1X (Figs 1B,C and S2B).The number of NPs inside the cells increased with time; after 20 h, it was higher with PS-COOH than with PS-NH2 at 0.01 and 1X concentrations (Figs 1B,C and S2B-D). The particles formed aggregates associated with WGA-labeled structures, whereas particles lying in cell-free areas were not labeled (S3A,B Fig). Cells located at the periphery of the outgrowth, which probably emerged first from the neural tube and were therefore exposed for longer, appeared to contain more NPs (Fig 1A). Cultures exposed to high concentration (10X) of NPs for 20 hours showed that NCCs could accumulate large quantities of particles, which were concentrated primarily near the nucleus, with in some cells an irregular nuclear shape in contact with these accumulations (Fig 1D, arrows). The PS-NPs persisted in cells, even when the medium had been replaced with the control medium for an additional day, and the cells remained positive for the Sox-10 (S3C Fig), a marker of migratory NCCs which plays a critical role in the maintenance of their progenitor state (43).

**Figure 1:**
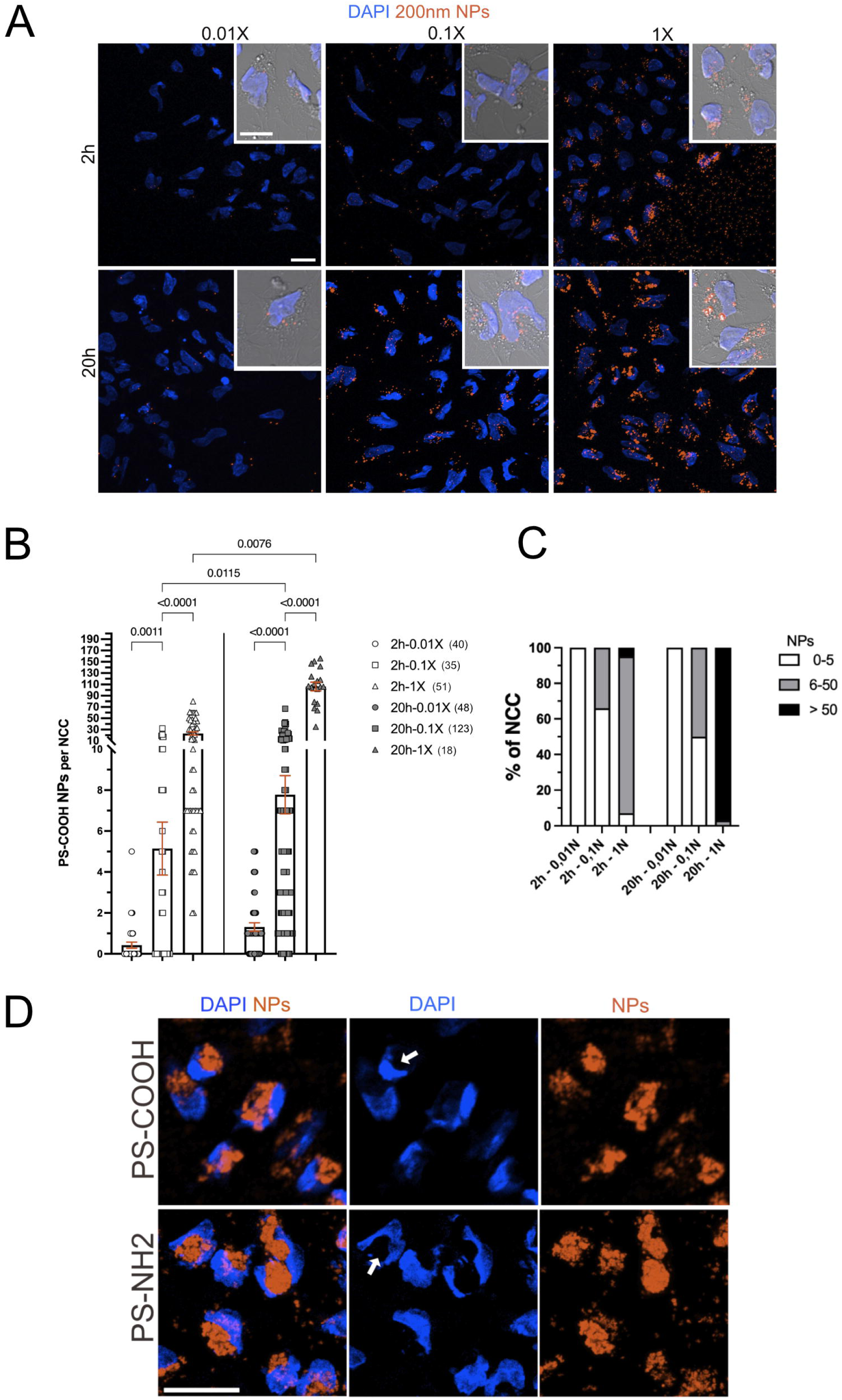
Dose and time-dependent incorporation of carboxylate- and amine-functionalized 200nm-PS NPs in NCCs. (A) Z-projection of confocal acquisition images of NCCs after 2 hours (h) (top panels) and after 20 h (bottom panels) of exposure to PS-COOH NPs (red) at the indicated concentrations (0.01X, 0.1X and 1X) and with DAPI visualization of the nuclei (blue). The inserts show a merge image of phase contrast, DAPI and NPs at higher magnification. (B) Scatter plots of the number of PS-COOH NPs incorporated per cells after 2 h (white bars) and after 20 h (grey bars) of exposure at the indicated concentrations. One dot per cell. Error bars are s.e.m. (C) Proportion of NCC having incorporated 0-5 NPs (white bars), 6-50 NPs (grey bars) or more than 50 PS-COOH NPs (black bars) after 2h and after 20 h of exposure, calculated from data in B. (D) Z-projection of confocal acquisition images of NCCs after 20 h of exposure to PS-COOH or PS-NH2 NPs (red) at 10X and with DAPI visualization of the nuclei (blue). The arrows point to nuclei of irregular shape in NCCs containing large accumulations of NPs. (n) in B indicate the number of NCC analyzed per condition. Scale bars in the panels and inserts in A are 25 and 50 μm, respectively. Scale bar in D, 20 μm. Data are from two independent experiments. In A and D, images are representatives from these experiments (n = 3 explants). Data in B were compared between the different concentrations at each time point and between the two type points at each concentration and analyzed using non-parametric Kruskal-Wallis with uncorrected Dunn’s test, with only the significantly different P values <0.05 indicated; comparisons not indicated are not significant.

These results indicate that when exposed to PS-NPs, NCCs can rapidly incorporate high numbers of particles without overt cell loss or detachment and that the surface functionalization had a moderate effect on their uptake.

### Polymer type and surface functionalization of nanoplastics are associated with distinct alterations of NCC outgrowth, density and spreading

Using our culture system, in which NCCs can rapidly internalize PS NPs in numbers ranging from a few to nearly a hundred per cell within 2 hours of exposure to 1X (Fig 1 and S2 Fig), we analyzed the effects of NP exposure on NCC behavior. We first analyzed the effects of NPs on NCC dispersion. Under control condition, NCCs delaminated from the neural tube and migrated collectively to rapidly produced large and dense outgrowth of cells during the first 24 hours of culture and with well-spread NCC morphology (Fig 2A). Similarly, in the presence of PS-COOH or PS-NH2, the neural tube rapidly produced large and dense outgrowths of NCCs up to 1X concentration (Fig 2A). 10X concentration of PS-COOH or PS-NH2 provoked a moderate but significant reduction in the outgrowth area (Fig 2B, left). In the presence of PS-COOH at 10X, the reduced NCC outgrowth area was accompanied by an increase in cell density (Fig 2B, right), whereas no difference was detected for PS-NH2. In addition, PS-COOH and PS-NH2 decreased NCC spreading from 0.01X, with a further decrease as the dose increased (Fig 2C).

**Figure 2:**
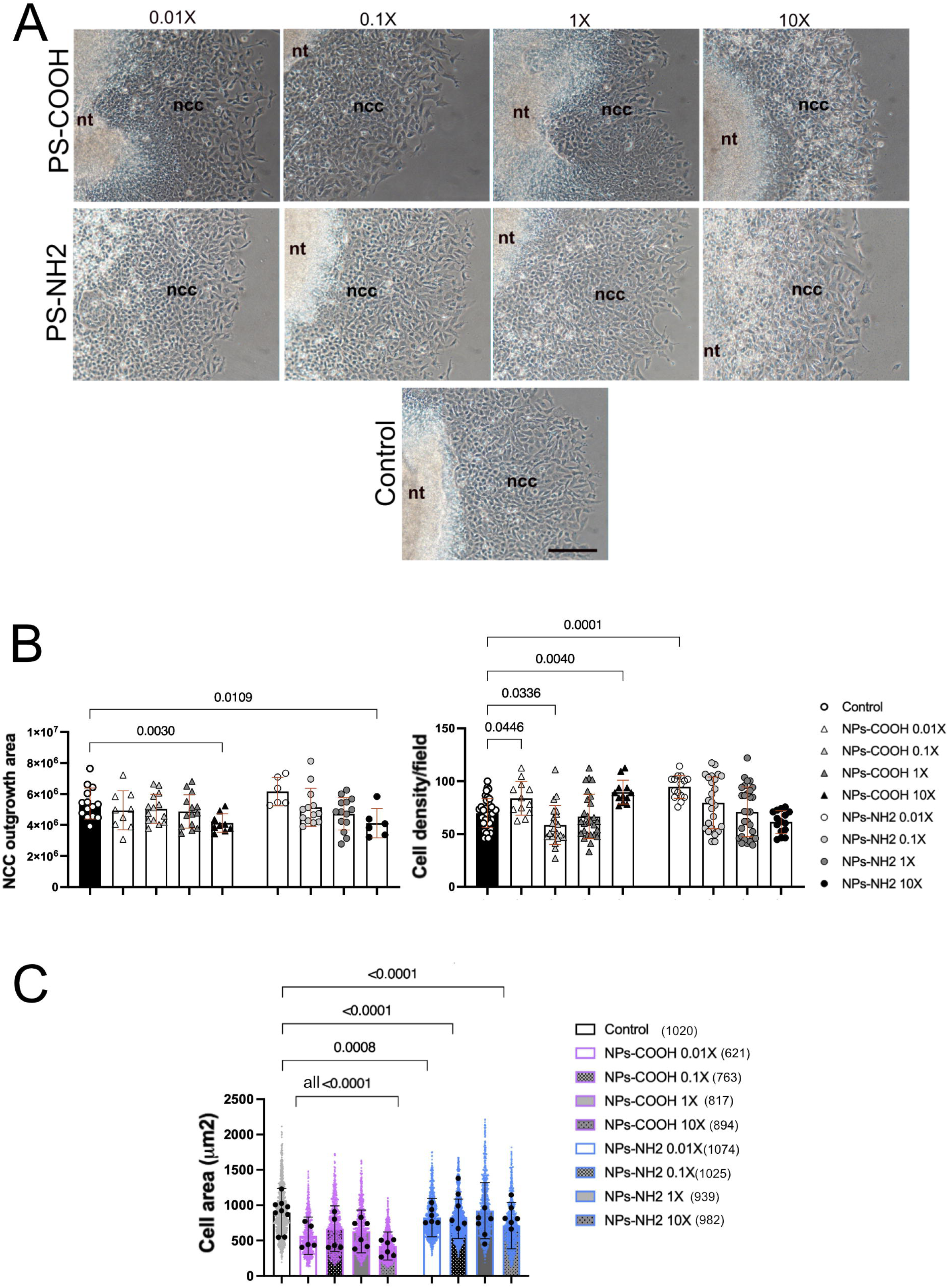
Dose-dependent effect of carboxylate- and amine-functionalized 200nm-PS NPs on NCC outgrowth, density and spreading. (A) Phase contrast images of NCC outgrowths after 24 h of exposure to PS-NPs at the indicated concentrations or in control condition (without NPs). nt: neural tube, ncc: neural crest cells. Scale bar, 100 μm. (B) Left panel: Scatter plot of NCC outgrowth area in μm^2^ produced after 24 h of exposure to PS-NPs at 0.01X, 0.1X, 1X indicated concentrations or in control condition (one dot per outgrowth; n = 9, 14, 14 and 9 for PS-COOH; n = 6, 13, 14 and 6 for PS-NH2 at 0.01X, 0.1X, 1X and 10X, respectively; n = 14 for control). Right panel: Scatter plot of NCC density in the same culture conditions as in B and for each outgrowth shown in the left panel in B. One dot represents the quantification in each image taken to cover each outgrowth; n = 12, 22, 27 and 12 for PS-COOH; n = 15, 25, 30 and 15 for PS-NH2 at 0.01X, 0.1X, 1X and 10X, respectively; n = 37 for control). (C) Superplot of the NCC spreading (cell area) after 24h of exposure to PS-COOH (pink bars), PS-NH2 (blue bars) NPs at 0.01X, 0.1X, 1X and 10X or in control condition (black bars). Values for individual cells are shown as transparent dots and the mean of each explant as a black dot. Bars indicate the mean ± S.D. calculated on all cells. Number of explants analyzed; n = 5, 5, 6 and 6 for PS-COOH and n = 6, 6, 6 and 6 for PS-NH2 at 0.01X, 0.1X, 1X and 10X, respectively, and 9 for control. (n) in C indicate the total number of NCCs analyzed per condition. Data are from at least three independent experiments. Images in A are representatives from these experiments. Error bars in B and C are S.D. Data in B and C were compared relative to the control and analyzed using non-parametric Kruskal-Wallis with uncorrected Dunn’s test, the tests in C being performed on cell values, with only the significantly different P values <0.05 indicated; comparisons not indicated are not significant.

We next assessed the effects of the other plastic polymer types and the surface functionalization on NCC dispersion. In contrast to PS-NH2, both PP-NH2 (Figs 3A-B) and PE-NH2 (Figs 3C-D) greatly affected neural tube explant cultures in a dose-dependent manner. We noticed more than three-fold reduction of NCC outgrowth area at 1X, compared to control, with more round or elongated NCCs with a tendency to aggregate, increasing locally the cell density (Figs 3A-D and S4B,C). At 10X concentration, only a small amount of NCCs remained, and the neural tubes had detached from the substrate in the presence of PP-NH2 and PE-NH2 (Figs 3A,C and S4B,C), a phenomenon that was not observed in the presence of PS-NPs (Figs 2A, S4A). PP- and PE-COOH reduced NCC outgrowth less than their NH2 counterparts (Fig 3A-D) whilst no difference was detected with the non-functionalized PP up to 1X (Fig 3B, upper panel). In addition, all forms of PP- and PE-NPs decreased cell spreading from 0.01X concentration, in a dose-dependent manner. The negative effect of NPs on cell spreading appeared more pronounced for amine-functionalized forms of PP and PE compared to carboxylate-functionalized forms and non-functionalized PP (Fig 3B,D, lower panels).

**Figure 3:**
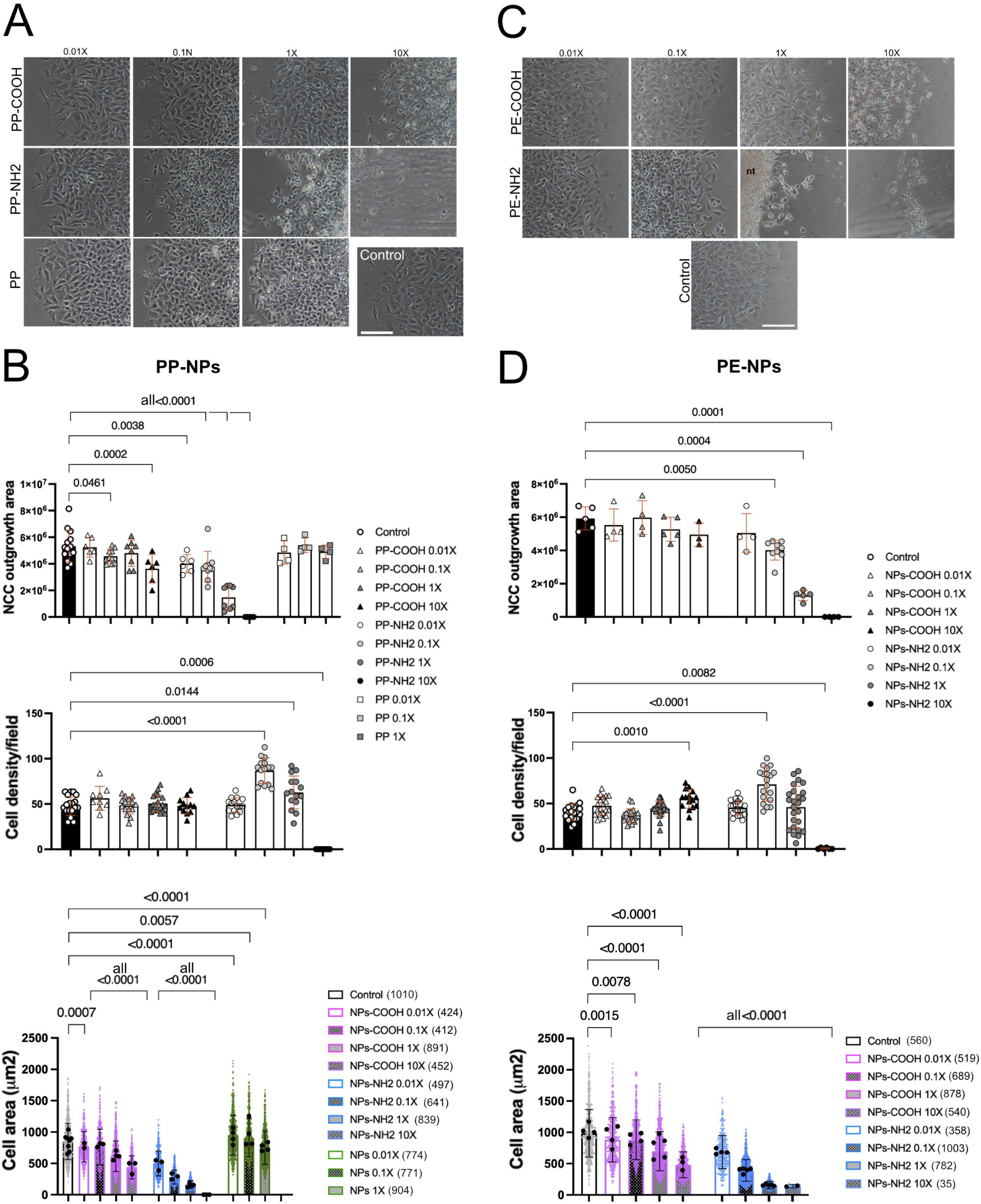
Dose-dependent effect of carboxylate- and amine-functionalized 200nm-PP and PE NPs on NCC outgrowth, density ad spreading. (A) Phase contrast images of NCC outgrowths after 24 h of exposure to PP-COOH, PP-NH2 and PP NPs at the indicated concentrations or in control condition (without NPs). (B) Top panel: Scatter plot of NCC outgrowth area produced in 24 h cultures in the same culture conditions as in A (one dot per outgrowth; n = 6, 10, 8 and 6 for PP-COOH, and n = 6, 10, 8 and 6, for PP-NH2 at 0.01X, 0.1X, 1X and 10X, respectively; n = 4, 4 and 4 for PP at 0.01X, 0.1X, and 1X, respectively; n = 14 for control). Explants detached from the substratum, with no outgrowth, were assigned an area of 0. Middle panel: Scatter plot of NCC density in the same culture conditions as in A and for each outgrowth shown in B, top panel. Explants detached from the substratum, with no outgrowth, were assigned a density of 0. One dot represents the quantification in each image taken to cover each outgrowth; n = 9, 15, 20 and 12 for PP-COOH and n = 15, 15, 15 and 10 for PP-NH2, at 0.01X, 0.1X, 1X and 10X, respectively; n = 20 for control). Density was not quantified for cultures exposed to PP. Bottom panel: Superplot of the NCC spreading (cell area) in 24 h explants in the same culture conditions as in A with PP-COOH (pink bars), PP-NH2 (blue bars), PP NPs (green bars) or in control condition (black bars). Values for individual cells are shown as transparent dots and the mean of each explant as a black dot. Bars indicate the mean ± S.D. calculated on all cells. Number of explants analyzed are n = 2, 3, 4 and 3 for PP-COOH and n = 3, 4, 4 and 3 for PP-NH2 at 0.01X, 0.1X, 1X and 10X, respectively; n = 4, 4, and 4 for PP at 0.01X, 0.1X and 1X, respectively; n = 6 for control. (C) Phase contrast images of NCC outgrowths after 24 h of exposure to PE-COOH and PE-NH2 NPs at the same concentration as in A or in control condition. nt, neural tube. (D) Top panel: Scatter plot of NCC outgrowth area produced in 24 h cultures in the presence of NPs or in control condition (one dot per outgrowth; n = 4, 4, 5 and 3 for PE-COOH, n = 4, 9, 5 and 4 for PE-NH2 at 0.01X, 0.1X, 1X and 10X, respectively, and 5 for control). Explants detached from the substratum, with no outgrowth, were assigned an area of 0. Middle panel: Scatter plot of NCC density in the same culture conditions as in C and for each outgrowth shown in D, top panel. Explants detached from the substratum, with no outgrowth, were assigned a density of 0. One dot represents the quantification in each image taken to cover each outgrowth; n = 20, 20, 25 and 15, respectively for PE-COOH, n = 20, 19, 25 and 6, respectively for PE-NH2, and 20 for control). Bottom panel: Superplot of the NCC spreading in 24 h primary cultures in the same culture conditions as in C with PE-COOH (pink bars), PE-NH2 (blue bars) or in control condition (black bars). Some explants detached from the substratum, with no outgrowth had few sparse NCCs from which cell spreading was calculated. Values for individual cells are shown as transparent dots and the mean of each explant as a black dot. Bars indicate the mean ± S.D. calculated on all cells. Number of explants analyzed are n = 4, 4, 5 and 4 for PE-COOH and n = 4, 5, 5 and 2 for PE-NH2, at 0.01X, 0.1X, 1X and 10X, respectively; n = 4 for control). Scale bar in A and C, 100 μm. (n) in B and D, bottom panels, indicate the total number of NCCs analyzed per condition. Error bars in B and D are S.D. Data are from at least three independent experiments. Images in A and C are representatives from these experiments. Data were analyzed using non-parametric Kruskal-Wallis with uncorrected Dunn’s test, the tests in the bottom panels being performed on cell values, with only the significantly different P values <0.05, all relative to control indicated, the comparisons not indicated are not significant. In B and D, bottom panels, data on cell values were also analyzed using non-parametric Kruskal-Wallis with uncorrected Dunn’s test to compare between functionalized forms of PP and PE NPs, the significantly different P values were all <0.0001 but are not indicated on the Superplots for clarity.

These findings indicate that NCC outgrowth differed according to the polymer type and the surface functionalization of the NPs they were exposed to. PP and PE have a more detrimental effect on NCC outgrowth than PS, particularly when functionalized with amine groups. In addition, exposure to each type of NPs altered NCC shape and spreading on the substrate.

### Effects of polymer type and surface functionalization of nanoplastics on NCC proliferation and apoptosis

NCCs are highly proliferative progenitor cells during their collective migration, a process that contributes to rapid dispersion and colonization of their environment in primary culture (37), in a timing similar to what is observed in vivo (30, 44). Thus, we investigated the impact of NPs on the expression of Sox-10, a marker of migratory and multipotent NCCs (37, 43), and on NCC proliferation. Almost all NCCs found in the outgrowths at 24 hours, in the presence of both -COOH or -NH2 forms of PS, PP and PE NPs up to 1X concentrations, expressed Sox-10 (Fig 4A). Compared to control condition, the presence of NPs affected NCC proliferation with increasing doses, but with different severity regarding the polymer functionalization and type. NCC proliferation was not decreased by NPs at lower doses. PS-NH2 moderately affected cell proliferation, whilst PP- and PE-NH2 decreased it by more than two-fold at 1X concentration (Fig 4B-E). At 10X concentration, PS-NH2-treated cultures still presented 30% of proliferative NCCs in the outgrowths whilst in PP- and PE-NH2-treated cultures the rare NCCs found were all EdU negative (Fig 4C-E), in line with the loss of outgrowth observed at 24 hours (Fig 3A,C). PS-COOH also decreased proliferation: the decrease was visible from 0.01X to 0.1X, then stabilized, and was statistically significant at 1X and 10X (Fig 4B). In contrast to PS, the impact of PP- and PE-COOH on proliferation was apparently less pronounced than their NH2 counterparts (Fig 4B-E).

**Figure 4:**
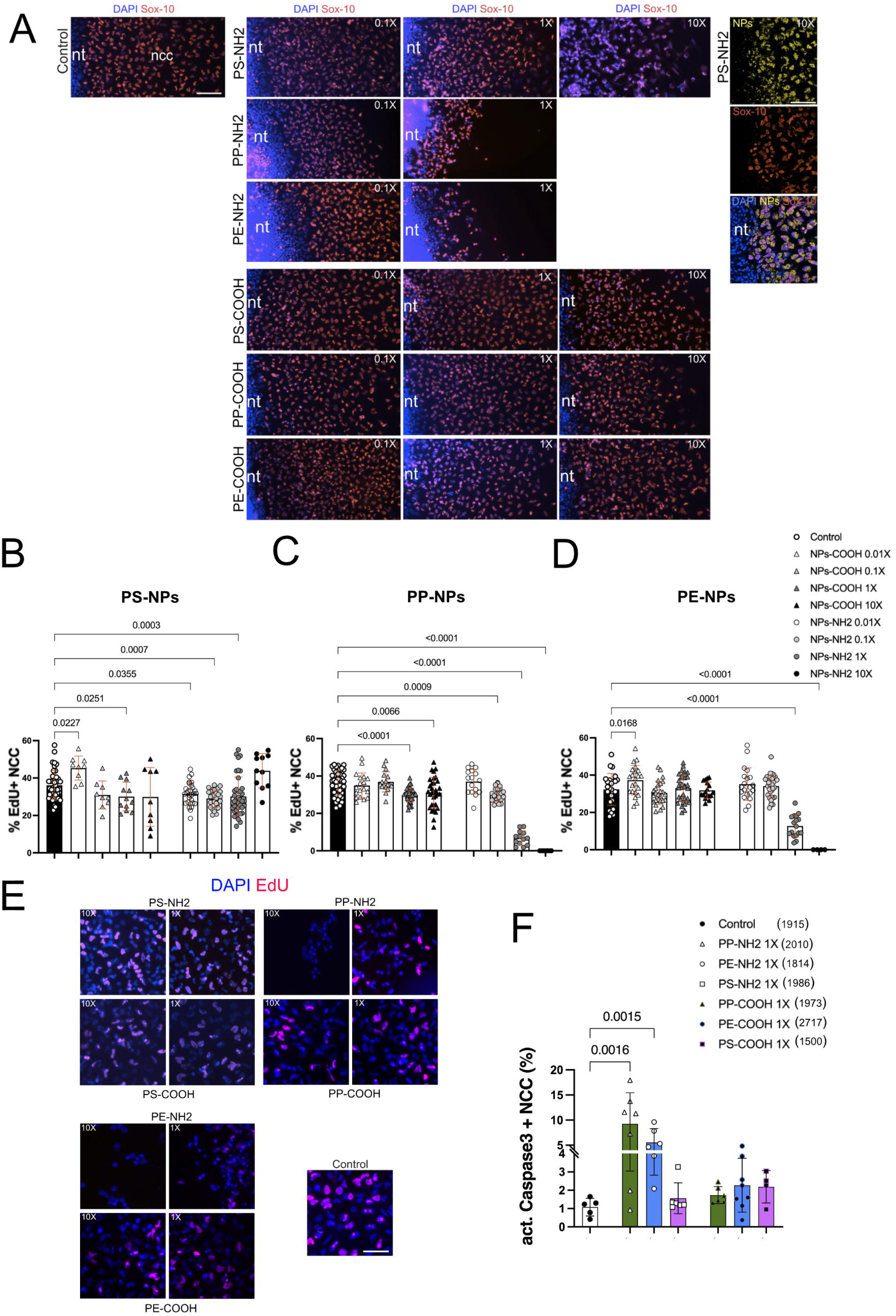
Effects of carboxylate- and amine-functionalized 200nm NPs on Sox-10 expression, proliferation and apoptosis of NCCs. (A) Merge Images for Sox-10 staining (NCCs, red) and nuclei (DAPI, blue) on NCC cultures after 24 h exposure to NPs-NH2 and - COOH at the indicated concentrations, and in control condition (without NPs). Images for PP-NH2 and PE-NH2 at 10X are not shown because the neural tubes had detached at 24 h. nt: neural tube. Scale Bar, 100 μm. The images in the far-right column show the highest magnification of the PS-NH2 NPs distribution (yellow), Sox-10 staining (red), and nuclei (blue) in NCC cultured at 10X concentration. nt, neural tube; ncc, neural crest cells. Scale bar, 50 μm. (B) Scatter plots of the % of EdU positive NCCs in 20 h cultures (including 1 h of EdU incorporation) in the presence of PS-NPs at 0.01X, 0.1X, 1X and 10X concentrations, and in control condition with mean ± S.D. One dot represents the quantification from each image (n = 9, 9, 12 and 9 for PS-COOH; n = 29, 29, 40 and 11 for PS-NH2 at 0.01X, 0.1X, 1X and 10X, respectively, and n = 42 for control) taken to cover as much as possible of the area occupied by the NCC population around the neural tube for different explants with n explants = 3, 3, 4 and 4 for PS-COOH; n = 3, 3, 7 and 4 for PS-NH2 at 0.01X, 0.1X, 1X and 10X, respectively, and n = 7 for control. The number of NCCs counted is 1819, 2185, 3699, and 1860 for PS-COOH; 10915, 11318, 14101, and 5960 for PS-NH2 at 0.01X, 0.1X, 1X and 10X, respectively; 18975 for control. (C) Scatter plots of the % of EdU positive NCC in 20 h primary cultures in the presence of PP-NPs at 0.01X, 0.1X, 1X and 10X, and in control conditions, with mean ± S.D. One dot represents the quantification from each image (n = 18, 17, 36 and 32 for PP-COOH; n = 16, 23, 13 and 7 for PP-NH2 at 0.01X, 0.1X, 1X and 10X, respectively, and n = 45 for control) taken to cover as much as possible of the area occupied by the NCC population around the neural tube for different explants with n explants = 3, 3, 5 and 6 for PP-COOH; n = 3, 3, 5 and 5 for PP-NH2 at 0.01X, 0.1X, 1X and 10X, respectively and n = 9 for control. The number of cells counted is 9941, 5428, 12283, and 11943 for PP-COOH; 7119, 8186, 3350 and 94 for PP-NH2 at 0.01X, 0.1X, 1X and 10X respectively; 14258 for control. In PP-NH2 explants detached from the substratum, with few sparse NCCs. (D) Scatter plots of the % of EdU positive NCCs in 20 h primary cultures in the presence of PE-NPs at 0.01X, 0.1X, 1X and 10X, and in control conditions, with mean ± S.D. One dot represents the quantification from each image (n = 22, 26, 34 and 17 for PE-COOH; n = 20, 33, 17 and 4 for PE-NH2 at 0.01X, 0.1X, 1X and 10X, respectively, and 25 for control) taken to cover as much as possible of the area occupied by the NCC population around the neural tube for different explants with n explants = 4, 4, 5 and 3 for PE-COOH; n = 4, 5, 5 and 3 for PE-NH2 at 0.01X, 0.1X, 1X and 10X, respectively, and n = 4 for control. The number of cells counted is 7608, 7900, 8227, and 10149 for PE-COOH; 6599, 14718, 3848 and 85 for PE-NH2 at 0.01X, 0.1X, 1X and 10X, respectively; and 10053 for control. In PE-NH2 explants detached from the substratum, with few sparse NCCs. (E) Images of EdU incorporation (visualized in magenta) and DAPI-labelled nuclei (blue) of NCCs in 20 h of culture (including 1 h of EdU incorporation) in the presence of NH2- and COOH-PS, PP, PE-NPs, at 1X and 10X concentrations, and in the control condition (without NPs), Scale bar, 40 μm. (F) Scatter plots of the % of activated Caspase-3 positive NCCs in 20 h primary cultures in the control medium or in the presence of COOH- and NH2-PP (green bars), PE (blue bars), and PS (Pink bars) NPs at 1X, with mean ± S.D. One dot represents the quantification from each image analyzed (n = 6, 7 and 6, respectively for PP-, PE-, and PS-NH2; n = 8, 6, and 4, respectively for PP-, PE-, and PS-COOH, and n = 5 for control) taken on each side of the neural tube when NCCs had emigrated on both sides, and on one side only otherwise, to cover as much as possible of the NCC population, for different explants (n = 2 for control and NPs-NH2; 2 for PS-COOH, and 3 for PP- and PE-COOH). (n) indicate the total number of NCCs analyzed per condition. Data are from two independent experiments. Images in A and E are representatives from these experiments. Data in B and C were analyzed using the non-parametric Kruskal-Wallis test followed by Dunn’s test, data in D using one-way ANOVA followed by Fisher’s least significant difference (LSD) test, and data in F using the Kruskal-Wallis test followed by Dunn’s test, all relative to the control, without correction for multiple comparisons, with only the significantly different P values <0.05 indicated; comparisons not indicated are not significant.

The NCC apoptosis was evaluated in cultures exposed at 1X concentration of NPs using anti-activated caspase3 antibody. In control condition, very few NCCs underwent apoptosis during their dispersion from neural tube explant after 20 hours of culture, with 1% of cells expressing activated caspase 3. In contrast, PP- and PE-NH2 induced a significant increase in number of apoptotic NCCs, about nine-fold for PP-NH2 and five-fold for PE-NH2, compared to control, whilst no difference from control was detected with PS-NH2 and COOH-NPs (Fig 4F).

Our results indicate that the effects of the NPs on NCC proliferation and apoptosis differed according to the polymer type tested and surface functionalization. These parameters should be considered when evaluating the impact of NPs on cells. PP and PE had a stronger effect than PS on NCC proliferation and apoptosis, and particularly with NH2-functionalization. NCCs remained Sox-10-positive in all conditions analyzed.

### Effects of nanoplastics on NCC migration and production period

We analyzed the kinetics of the NCC outgrowth progression and cell migration properties using video-microscopy analyses. Neural tube explant cultures were performed in the control medium or in the presence of NH2-NPs, at 1X and 0.1X concentrations. In the control condition, a sustained progression of the NCC migration front was observed, traveling 300 μm in 12 hours (Fig 5, left). In the presence of PS-NH2 NPs at 1X concentration, the migration front advanced steadily but slowly, traveling 250 μm in 12 hours. In contrast, PP- and PE-NH2 induced a severe inhibition on the progression of NCCs migratory front, which rapidly slowed until it came to a halt after 7 hours of exposure to the NPs, traveling only 100 μm in 12 hours (Fig 5A, left). The average NCC speed in the presence of PP- and PE-NH2 at 1X was three times lower (Fig 5B, left), and cell persistence was increased (Fig 5B, right), compared to those measured in the presence of PS-NH2 or in control (Fig 5B). The impact of PP- and PE-NH2 NPs on front progression and migration parameters, at the 1X concentration, were evident within the first hour after the NPs deposition (Fig 5A,C). Compared to control, no difference was detected in the front’s progression of NCCs in the presence of each of the three NH2-NPs at 0.1X concentration (Fig 5A, right). No difference was detected in NCC speed and persistence with PP- and PE-NH2 NPs, whereas with PS-NH2, a slight reduction in cell speed and increase in their persistence was observed (Fig 5B).

**Figure 5:**
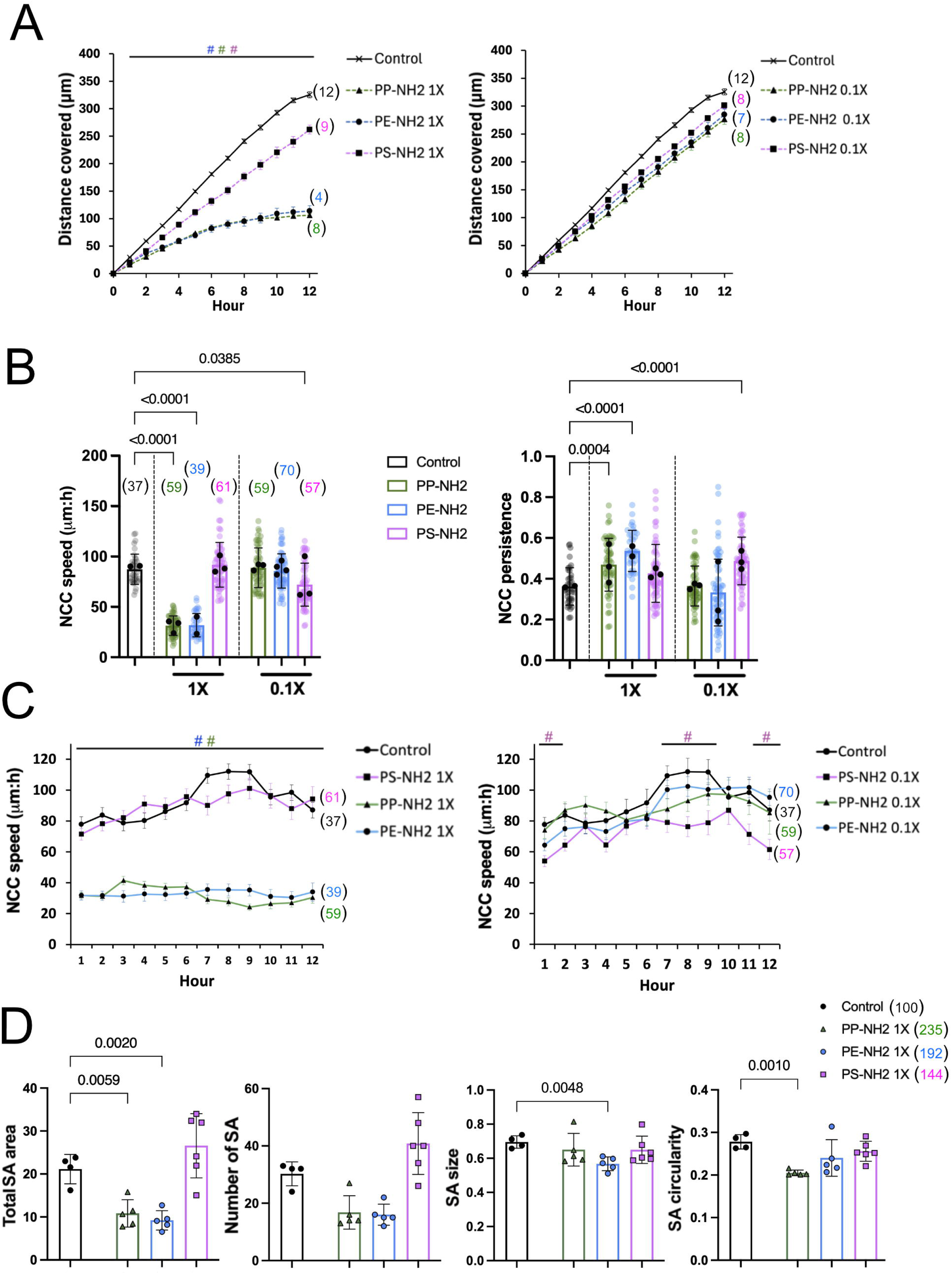
Effect of amine-functionalized 200nm NPs on NCC migration and substrate adhesion parameters. (A) Distance covered over time by the NCC migratory front, and measured using video-microscopy, in cultures exposed to PS- (magenta), PP- (green) and PE- (blue) NH2 NPs at 1X (left) and 0.1X (right) concentrations, and in control condition (without NPs, black; shared controls for NH2- and COOH-NPs conditions shown in Fig 5A and 6A). t0 corresponds to onset of recording, 3 h after initiation of culture, with the NPs applied 60 to 90 min before recording. Front progression was analyzed using a custom GUI written in Matlab. Values are mean ± s.e.m. (n) indicate the number of explants. (B) Superplots of the speed (left panel) and of the persistence (right panel) of individual NCCs at the indicated concentrations. Values for individual cells are shown as transparent dots and the mean of each explant as a black dot ± S.D, (n) indicate the total number of cells analyzed in the explants (n explants = 3, 2, 3 for PP-, PE- and PS-NH2 NPs at 1X, and 3, 4 and 3 for PP-, PE- and PS-NH2 at 0.1X, respectively; and 3 for control). Data were obtained throughout video-microscopy recordings and analyzed using MetaMorph software. (C) Graphs showing the hourly speed of NCCs in cultures exposed to PS- (magenta), PP- (green) and PE- (blue) NH2 NPs at 1X (left) and 0.1X (right) concentrations, and in control condition (without NPs, black), over a 12-hour period. Values are mean ± s.e.m. (n) indicate the total number of NCC analyzed in primary cultures (n explants as in B). (D) From left to right: Scatter plot of total substrate adhesion (SA) area and SA number per NCC, SA size and circularity, respectively, with mean ± S.D measured in 24 h cultures exposed to PS- (magenta), PP- (green) and PE- (blue) NH2 NPs at 1X concentration, and in control (without NPs, black). Each dot represents the quantification performed on one image (n images = 5, 5 and 6, respectively PP-, PE- and PS-NH2 NPs, respectively; n = 4 for control) taken from n explants, with n = 4, 4, 3 for PP-, PE- and PS-NH2 NPs 1X conditions, respectively; and 2 for control. (n) indicate the total number of NCCs analyzed per condition. Data in A-D are from two independent experiments. Data in A and C were analyzed at each hour in the presence of NPs by unpaired t-test two-tailed test relative to the corresponding time point of the control condition, with colored # for the P values <0.05 at the corresponding timepoints and conditions; comparisons not indicated are not significant. Data in B were analyzed using non-parametric Kruskal-Wallis with uncorrected Dunn’s test relative to the control, and data in D by one-way ANOVA followed by Fisher’s least significant difference (LSD) test for comparisons relative to the control; comparisons not indicated are not significant.

In the presence of PP- and PE-COOH NPs the advance of NCC migration front was slowed, but to a lesser extent than with their NH2-counterparts. With PP- and PE-COOH at 1X concentration, the migration front advanced steadily but more slowly than in the control condition, traveling 230 and 290 μm, respectively, in 12 hours, compared to 330 μm for the control (Fig 6A, left). The effect of PP-COOH became apparent within the first few hours of exposure, whereas it was only significantly visible after 9 hours for PE-COOH (Fig 6A, left). At 0.1X concentration, a significant reduction in front progression was observed in the presence of PP-COOH, occurring after 7 hours of exposure to NPs, traveling 250 μm in 12 hours, while with PE-COOH no difference in the front progression was observed, with 300 μm covered in 12 hours (Fig 6A, right). In the presence of PS-COOH NPs at the two doses tested, no difference in front progression was observed, compared to control (Fig 6A). Compared to the control, the average NCC speed was reduced by one-third in the presence of PP- and PE-COOH at 1X concentration, whereas it was almost double in the presence of PS-COOH (Fig 6B, left). In addition, NCC persistence was decreased in the presence of PP- and PS-COOH at 1X (Fig 6B, right). In contrast, at 0.1X, all three COOH-NPs increased the average NCC speed (Fig 6B, left). The effects of COOH-NPs on NCC speed emerged within the first hour after NPs deposition (Fig 6C).

**Figure 6:**
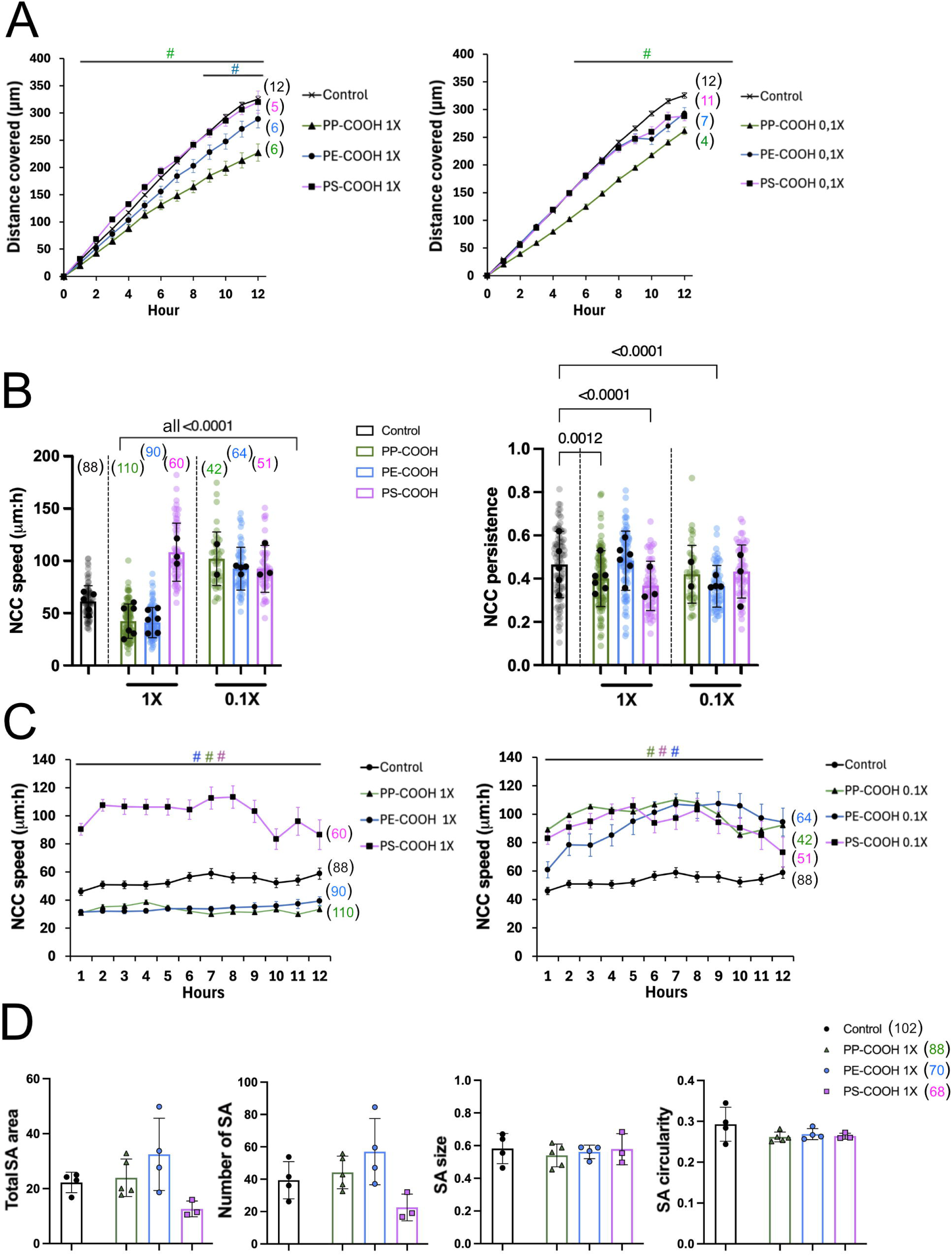
Effect of carboxylate-functionalized 200nm NPs on NCC migration and substrate adhesion parameters. (A) Distance covered over time by the NCC migratory front in cultures exposed to PS (magenta), PP (green) and PE (blue) NPs-COOH at 1X (left) and 0.1X (right) concentrations, and in control condition (without NPs, black; shared controls for NH2- and COOH-NPs conditions shown in Fig 5A and 6A), and measured and analyzed as described in Fig 5. Values are mean ± s.e.m. (n) indicate the number of explants. (B) Superplots of the speed (left panel) and of the persistence (right panel) of individual NCCs at the indicated concentrations. Values for individual cells are shown as transparent dots and the mean of each explant as a black dot ± S.D (n) indicate the total number of cells analyzed for the explants (for n explants = 6, 6, and 3 for PP-, PE- and PS-COOH NPs 1X, and = 2, 4 and 3 for in PP-, PE- and PS-COOH 0.1X, respectively; n = 5 for control). Data were obtained throughout video-microscopy recordings, analyzed using MetaMorph software (C) Graphs showing the hourly speed of NCCs in cultures exposed to PS- (magenta), PP- (green) and PE- (blue) COOH NPs at 1X(left) and 0.1X (right) concentrations, and in control condition (without NPs, black), over a 12-hour period. Values are mean ± s.e.m. and (n) indicate the total number of NCCs analyzed in primary cultures (n explants as in B), and as described in legend of Fig 5C. (D) From left to right: Scatter plot of total substrate adhesion (SA) area and SA number per NCC, SA size and circularity, respectively, with mean ± S.D measured in 24 h primary cultures exposed to PS- (magenta), PP- (green) and PE- (blue) COOH NPs at 1X concentration, and in control (without NPs, black). Each dot represents the quantification from one image (n = 5, 4 and 3, respectively in PP-, PE- and PS-COOH NPs conditions, respectively; n = 4 for control) taken from 2 explants for PP-, PE- and PS-COOH NPs at 1X, and for control. (n) indicate the total number of NCCs analyzed per condition. Data in A-D are from two independent experiments. Data in A and C were analyzed at each hour in the presence of NPs by unpaired t-test two-tailed test relative to the corresponding time point of the control condition, with colored # for the P values <0.05 at the corresponding timepoints and conditions; comparisons not indicated are not significant. Data in B were analyzed using non-parametric Kruskal-Wallis with uncorrected Dunn’s test relative to the control, and data in D by one-way ANOVA followed by Fisher’s least significant difference (LSD) test for comparisons relative to the control; comparisons not indicated are not significant.

Delamination, an epithelial-to-mesenchymal transition-associated process, is a key early event that enables newly generated NCC to detach from the dorsal neural tube, migrate and disperse away. We analyzed the impacts of NPs on the duration of NCC production from the neural tube. The NCC production time was reduced by half in the presence of PP-NH2 and PE-NH2 at the 1X concentration, compared to the control, whereas no difference was observed with PS-NH2 or the three NPs-COOHs at 1X, nor with the NPs at 0.1X concentration (S5A Fig).

Our results indicate that the formation of the NCC outgrowth, the progression of the migration front, and the migration parameters differ according to the type of polymer and the functionalization of the NPs. PP- and PE-NPs have stronger effects on NCC migration than PS-NPs, particularly those functionalized with amine groups compared to those functionalized with carboxylate groups. The impact on front progression may be related to alteration in cell speed, as well as to the persistence and duration of the NCC production period.

The video microscopy analyses also revealed that the morphological changes in NCC observed in the presence of PP- and PE-NH2, such as cell elongation, rounding, or a reduction in cellular protrusions (see Figs 3, S4), had occurred within less than 4 and 7 hours, of exposure to these NPs, respectively, and after more than 10 hours of exposure to their COOH counterparts (S5B Fig). In contrast, no clear morphological change was observed in NCCs in the presence of PS-NPs (S5B Fig).

### Effects of nanoplastics on NCC substrate adhesions

To gain insight into the impact of NPs on NCC substrate adhesion, we examined by immunolabeling the cellular localization of paxillin, a cytoplasmic protein of integrin-based adhesion complex, used as a marker of substrate adhesions (SA). NCCs developed SA both in control condition and in the presence of all NPs tested (S6A and S7 Figs) but to varying degrees. We thus quantified several parameters related to SAs, such as the total area of SA and the SA number per NCC, as well as SA size and circularity.

In 24 hours-cultures exposed to PP- and PE-NH2 at 1X concentration, a two-fold decrease in total area of SAs per NCC was observed, compared to control, along with a 25% decrease in SA circularity with PP-NH2, a 15% decrease in SA size with PE-NH2 (Fig 5D). We noticed a trend toward a decrease in the number of SAs per cell with PP- and PE-NH2, but not significant (p value=0.065 for both NPs). In contrast, no difference was detected after exposure to PS-NH2 or all the COOH-NPs at 1X (Figs 5D, 6D).

At the lower dose of 0.1X, no difference in SA parameters was detected in NCCs cultured with PP- and PE-NH2, whilst with PS-NH2 the total SA area per NCC was increased by 54% (S6B Fig). These results indicate that the effects on SAs differ according to the type of polymer, the functionalization of the NPs. PP- and PE-NH2 reduced SA area at 1X concentration but not at 0.1X, which may reflect alterations in SA formation or renewal only for upon PP- and PE-NH2 NPs accumulation.

Overall, our results show that PS, PP, and PE NPs rapidly act on NCCs to reduce their spreading, even when the cells have incorporated only a small number of NPs. When the cells have incorporated more NPs, they react differently depending on the type of polymer they are exposed to. PP and PE NPs, functionalized with amine groups, are more harmful than PS to interfere with numerous processes involved in regulating NCC substrate adhesion, migration, proliferation, and apoptosis, as well as NCC production period and their collective behavior. These processes are essential to ensure efficient NCC dispersion and the colonization of their environment. Thus, the presence and accumulation of those NPs may ultimately have an impact on the final fate of NCCs and their derivatives.

## Discussion

The behavior and fate of cells are strongly influenced by their environment, which determines their shape, adhesion, polarity and contractility, and regulates their proliferation, differentiation, and migration potential. Several environmental factors can act on cells, including pollutants such as NPs. A previous study performed on 2-days-primary cultures of NCCs exposed to 25nm PS-NPs revealed that these cells can rapidly internalize NPs (32). Moreover, after ex ovo injection of early avian embryos with 25, 60 or 900nm PS-NPs, aberrant neural tube morphogenesis and increased cell apoptosis at dorsal midline were observed with consequences on embryonic development and NCC-derived structures, in a dose and size-dependent manner (32, 33). In the present study we investigated the unexplored short-term effects of exposure to NPs on NCC adhesion, migration, proliferation and apoptosis, and more specifically the impact of three plastic polymers, PS, PP, and PE, and of carboxylate- and amine-surface functionalization on these processes. The three polymers used in this study have different densities, and the height of the medium varies from one culture support to another. Nevertheless, the fraction of the administered dose that reaches the cell layer within 24 hours, that is the delivered dose, should be of the same order of magnitude for all polymers and all supports, thereby making a comparison of their effects at equal administered doses meaningful. Under these conditions, we observed different responses from NCCs depending on the NP type and its surface modification.

We show that early migratory NCCs incorporate 200nm NPs in a couple of hours. The number of particles found in cells increases in a dose-dependent manner, and with time of exposure, with both amine- and carboxylate-functionalized NPs. NCCs could rapidly accumulate a high number of PS-NPs without overt cell loss or detachment. Exposure to PS NPs reduce NCC cell spreading. This reduction is already detectable at the lowest dose of PS-NPs tested (0.01X), a dose at which most cells contained only a few NPs, and it is also observed upon exposure to PP- and PE-NPs, for which internalization could not be measured directly. This reduction becomes more pronounced as the dose increases. Intracellular accumulation of NPs, demonstrated for PS only, probably also account for the PP- and PE NPs effects. PS-NPs incorporation moderately decreases the spreading, dispersion, and proliferation, of NCCs, without detectable effect on apoptosis. Interestingly, PS-NPs are not the particles that exert the most pronounced effects on NCCs. PP- and PE-NH2 produce a strong decrease in NCC proliferation than PS-NH2, and an increase in apoptosis, at 1X concentration. While the NP size and concentration, as well as culture conditions, are specific to each study, other cell types were shown to display specific sensitivity to PS and PE polymers (24, 45), with 200 nm and 300 nm PE-NPs being more deleterious to the viability of SH-SY5Y neuroblastoma cells (45) and human embryonic stem cells (24), respectively, than PS-NPs.

NPs bearing positive and negative surface charges were shown to have distinct toxicity and affect cell response differently. For example, Anguissola et al. (2014) showed that 50nm PS-NH2 lead to increased apoptosis in numerous cell types, including lung, endothelium, liver, kidney, macrophages, glia, and neuronal cells while plain and COOH-NPs had no effect (46). Increased toxicity of 100nm PS-NH2 was also demonstrated on macrophages (25). In our study, NCC responses differ according to polymer type and surface functionalization. PP- and PE-NH2 decrease NCC proliferation and migration, and increase apoptosis (revealed by increase activated caspase-3 staining) more strongly than their COOH or plain counterparts, whilst PS-NH2 and PS-COOH have weaker effects on proliferation and no detected effect on apoptosis.

The dispersion of NCCs around the neural tubes to form large outgrowths also differs depending on the nature of the polymer and the surface functionalization of the NPs: PP-NH2 and PE-NH2 inhibit this process more strongly than PS-NH2, while COOH-NPs have a weaker effect on this process. The decrease in NCC dispersion may result both from the negative effects of these PP-NH2 and PE-NH2 NPs on NCC proliferation and apoptosis, and from changes in their collective behavior accompanied with marked decrease in their migration speed. The altered migratory behavior is already apparent within the first hours of exposure. It most likely originates from direct effects on the main drivers of NCC motility, namely substrate adhesions, and the cytoskeleton, acting on their dynamics or on the associated signaling pathways. A contribution of transcriptional changes in migration-related pathways cannot be excluded at later time points but was not assessed in this study. It might also involve physical interactions between NPs and the cytoskeleton, in agreement with recent studies (28, 47). PS-NPs were shown to have adverse effects on cell migration and invasion (28, 47–49) or to stimulate it (50–52). In a microglial cell model, PP-NPs and PS-NPs produce distinct but opposite effect on cell migration with what we observe on NCCs, PP-NPs stimulating microglial chemotactic migration index while PS-NPs have no effect (53). This underscores the need to compare, within each single cellular model, the impact of NPs with different polymer compositions and surface functionalization to give an overview of NPs toxicity and impact on similar cell responses, avoiding extrapolation of the NPs effects observed from other model systems.

In addition to the cell response mediated by internalized NPs, several extracellular mechanisms may contribute. Gold nanoparticles were shown, by molecular simulations, to adsorb to fibronectin and alter its conformation (54). Titanium dioxide and silica nanoparticles were reported to impair fibronectin-mediated adhesion and spreading of pre-osteoblasts (55), and adhesion and migration of keratinocytes (56), respectively. Our substrates are coated with fibronectin, a direct interference of nanoplastics with the adhesive substrate may happen, but this effect and the impact of the polymer type remain to be investigated. NPs also adsorb proteins from biological fluids to form a protein corona, whose formation depends on particle characteristics, including surface functional groups (57). Corona formation may locally reduce the availability of adhesion-promoting proteins from the medium and may itself account for part of the differences observed between amine- and carboxylate-functionalized NPs. Finally, we defined 1X as the number of NPs required to form one close-packed monolayer over the culture surface. Even if every particle reached the surface, they would cover at most about 0.9% and 9% of it at 0.01X and 0.1X doses, respectively. Physical masking of the adhesive surface of the support by the NPs (55) is therefore unlikely to explain the effects observed at these doses, whereas it cannot be excluded at higher doses. Our data do not allow these extracellular contributions to be separated from the effects of internalized particles, and the two may coexist.

When neural tube explants developed outgrowths in culture, NCCs remain Sox-10 positive after 24 hours of exposure to NPs in all conditions analyzed. Time lapse recordings show that PP- and PE-NH2 exposure shortened the production period of NCCs from neural tube at 1X dose, whereas PS-NH2 or COOH-NPs have any effect. Likely explanations could be a decreased adhesion of delaminating NCCs, a process which we would have noticed by formation of non-adherent rows of cells at the neural tube border, but alternatively that delaminating or pre-migratory NCCs have stayed stuck at the interface between dorsal neural tube and culture surface. An increase in apoptosis or a reduction in proliferation, both observed in the outgrowths, may also occur in pre-migratory NCC. Another reason could be that exposure PP- or PE-NH2 alters pre-migratory NCC developmental gene programs. A next step in this study would be to determine whether exposing neural tubes or embryos (harvested before the delamination stage) to NPs can affect the NCC genetic regulatory networks (58), especially the epithelial-mesenchymal transition program, as has been observed during carcinogenesis (26, 59, 60).

NCCs can be cultured for up to one or two weeks and, under appropriate conditions, can either maintain their stem cell status or differentiate into various cell types, including neurons, glial cells, melanocytes, and myofibroblasts (61, 62). Given their extensive contribution to embryonic development and their ability to generate a broad range of cell types and tissues, alterations of NCC production, proliferation, apoptosis or migration may have important consequences for developmental processes. Whether the polymer type and the surface functionalization of NPs alter the multipotency and differentiation potential of NCCs in vitro, and affect NCC-dependent developmental processes in vivo, therefore remains an important question to address.

From a human health perspective, these questions are particularly relevant because NCCs give rise to numerous cell types and tissues, including craniofacial tissues, melanocytes, cardiac and neuroendocrine derivatives, and components of the peripheral and enteric nervous systems (30, 31). Consequently, alterations in NCC development can contribute to congenital abnormalities and developmental disorders (31, 63). Our findings that specific NPs can perturb NCC proliferation, apoptosis, adhesion and migration raise the possibility that NP exposure could interfere with developmental processes that depend on NCCs. However, how these cellular effects translate into alterations of NCCs-derived tissues in vivo, at relevant exposure levels, remains to be established.

Our study highlights that commercially available NPs have minimal to detrimental depending on the degree of intracellular accumulation, polymer type, and surface functionalization. However, the NPs encountered by living organisms have undergone multiple transformations during their use, recycling and exposure to the environment. They adsorb various pollutants or other compounds and are subjected to physical stresses that modify their shape, surface properties and roughness. These factors may act synergistically with polymer type and potentially exacerbate their deleterious effects on cells and organisms, possibly at lower doses. It will therefore be important to further investigate how particle shape and surface properties, as well as the adsorption of environmental pollutants, influence NCC behavior and developmental processes. This will require approaches capable of reproducing, at least in part, the physicochemical complexity of environmentally derived NPs.

## Acknowledgments

The authors thank Sophie Féréol for helpful discussions, and Alwyn Dady for comments on the manuscript. The authors thank Xavier Decrouy from the IMRB imaging platform.

## Data availability

All relevant data are within the manuscript and its Supporting Information file.

## Author contributions

Conceptualization: Sylvie Dufour.

Data curation: Sylvie Dufour. Redouane Fodil

Formal analysis: Sylvie Dufour, Redouane Fodil

Funding acquisition: Sylvie Dufour, Redouane Fodil, Fred Relaix.

Investigation: Sylvie Dufour

Methodology: Sylvie Dufour and Redouane Fodil

Project administration: Sylvie Dufour

Resources: Sylvie Dufour, Redouane Fodil, Fred Relaix

Software: Redouane Fodil

Supervision: Fred Relaix Validation: Sylvie Dufour

Visualization: Sylvie Dufour

Writing – original draft: Sylvie Dufour

Writing – review & editing: Sylvie Dufour, Redouane Fodil, Fred Relaix

## Supporting information

**S1 Fig. Strategy used in the study for investigating the impacts of 200nm-plastic particles (NPs) on NCC primary cultures.**

**S1 Appendix. Nanoplastic dosimetry**

**S2 Fig, related to Fig 1. Dose and time-dependent incorporation of amine-functionalized 200nm-PS NPs in NCC.**

**S3 Fig, related to Fig 1. NCC incorporated and accumulated 200nm-NPs.**

**S4 Fig, related to Figs 2 and 3. Effect of 200nm-NPs on cell morphology on NCC primary cultures.**

**S5 Fig, related to Figs 5 and 6. Effect of 200nm-NPs on NCC production time and spreading.**

**S6 Fig, related to Fig 5. Effect of 200nm-NPs-NH2 on NCC substrate adhesion.**

**S7 Fig, related to Fig 6. Effect of 200nm-NPs-COOH on NCC substrate adhesion.**

