## supplementary information for "Polymer type and surface functionalization determine the impact of nanoplastics on neural crest cell adhesion, migration, proliferation, and apoptosis in primary cultures"

### Supporting information

**S1 Fig. Strategy used in the study for investigating the impacts of 200nm-plastic particles (NPs) on NCC primary cultures.**

**S1 Appendix. Nanoplastic dosimetry**

**S2 Fig, related to Fig 1. Dose and time-dependent incorporation of amine-functionalized 200nm-PS NPs in NCC.**

**S3 Fig, related to Fig 1. NCC incorporated and accumulated 200nm-NPs.**

**S4 Fig, related to Figs 2 and 3. Effect of 200nm-NPs on cell morphology on NCC primary cultures.**

**S5 Fig, related to Figs 5 and 6. Effect of 200nm-NPs on NCC production time and spreading.**

**S6 Fig, related to Fig 5. Effect of 200nm-NPs-NH2 on NCC substrate adhesion.**

**S7 Fig, related to Fig 6. Effect of 200nm-NPs-COOH on NCC substrate adhesion.**

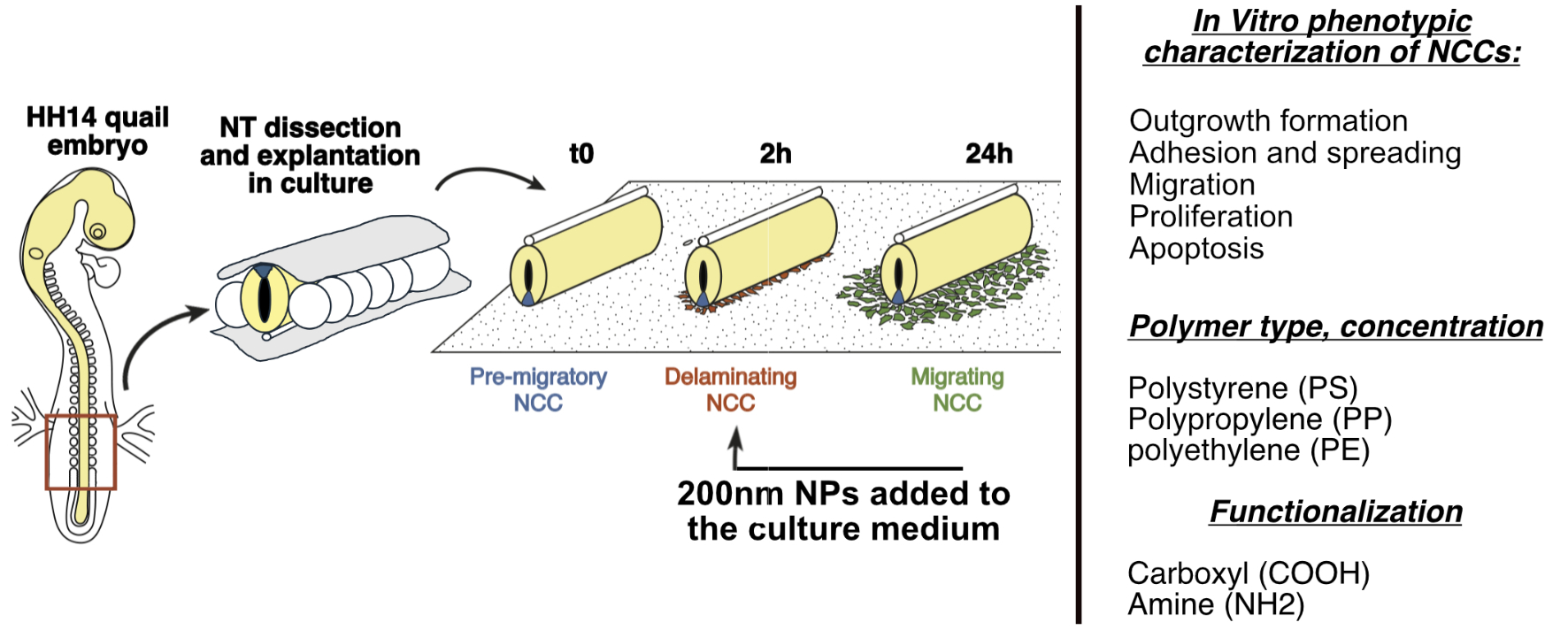

**S1 Fig. Strategy used in the study for investigating the impacts of 200nm-plastic particles (NPs) on NCC primary cultures.** Schematic representation of procedure of the neural tube explants culture at stages HH14 (see Methods for details), with indication of the timing in hour (h) of in vitro NCC outgrowth development around the neural tube and the arrow indicates the time of NPs deposition in the cultures. The embryonic region collected for neural tube preparation is delineated with a red rectangle.

**S1 Appendix. Nanoplastic dosimetry.** Equal mass concentrations of particles differing in density or composition do not deliver equal particle numbers or equal surface doses to adherent cells (1). Doses were therefore normalized to the culture surface rather than to the medium volume. One X was defined as the number of NPs forming a single hexagonally close-packed monolayer over the culture area:

$X=\frac{\phi A}{{\pi d^{2}}/4}$	(1)

where A is the growth area, d the nominal particle diameter (200 nm) and $\phi=\pi/\left( 2\sqrt{3} \right)=0.9069$ the two-dimensional close-packing fraction (2). This definition yields an identical administered dose per unit area on all three supports used, namely 2.88 × 10⁹ NPs cm⁻² at 1X (Table 1). Equivalent mass concentrations were calculated as:

$C=\frac{X\rho\pi d^{3}}{6V}$	(2)

where $\rho$ is the polymer density specified by the manufacturer (1.03, 0.89 and 0.88 g cm⁻³ for PS, PP and PE, respectively) and V the final volume of the culture medium.

Surface normalization equalizes the administered but not the dose delivered to the cell monolayer, since all three polymers are buoyant or nearly neutrally buoyant in aqueous culture medium (ρ ≈ 1.00 g cm⁻³).

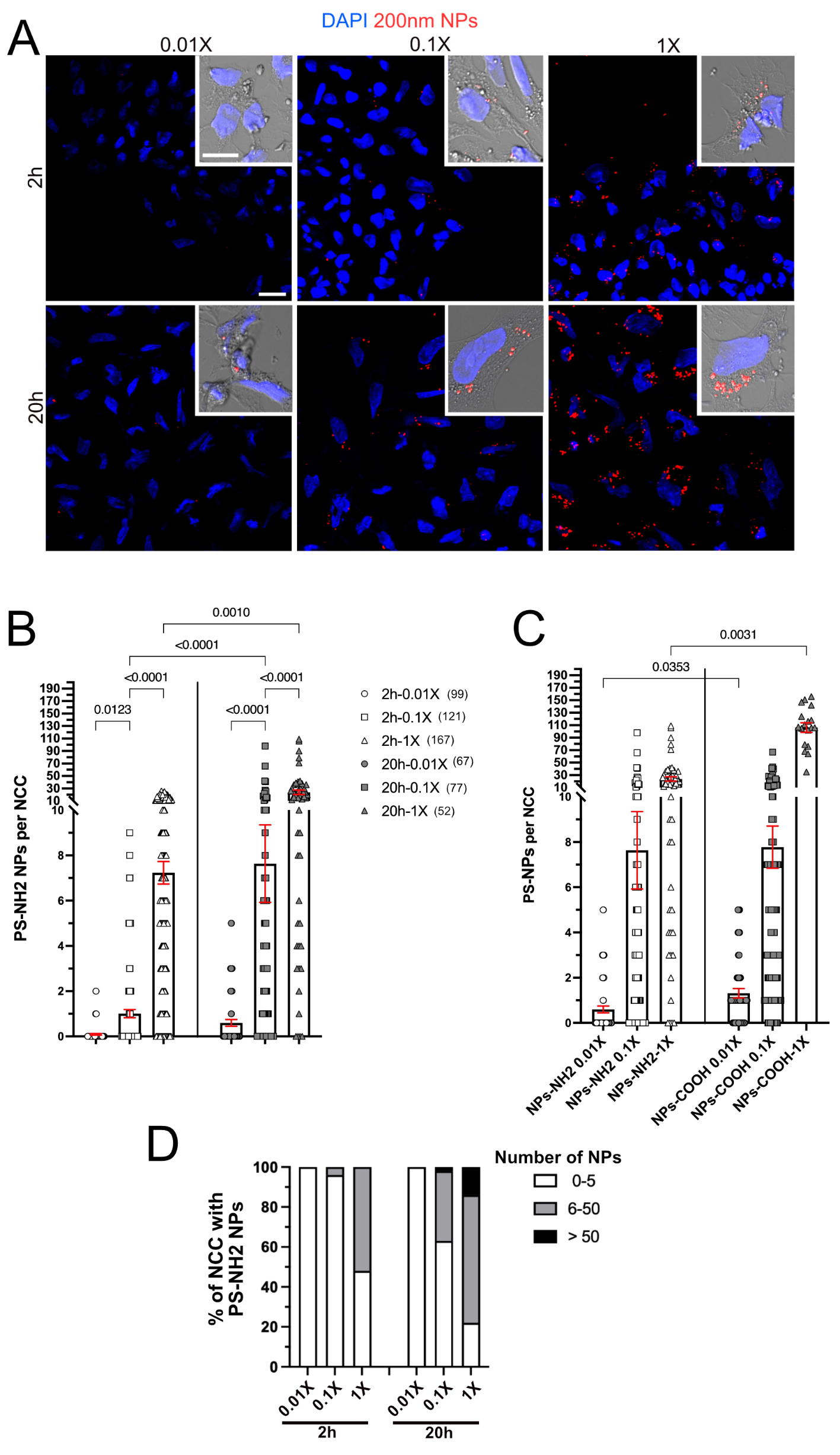

**S2 Fig, related to Fig 1. Dose and time-dependent incorporation of amine-functionalized 200nm-PS NPs in NCCs.** (A) Z-projection of confocal acquisition images of NCCs after 2 hours (h) (top panels) and after 20 h (h) (bottom panels) of exposure to PS-NH2 NPs (red) at the 0.01X, 0.1X and 1X concentrations, respectively, and with DAPI visualization of the nuclei (blue). The inserts show a merge image of phase contrast, DAPI and NPs at higher magnification. (B) Scatter plots of the number of PS-NH2 NPs incorporated per cells after 2 h and after 20 h of exposure at the same concentrations as in A. One dot per cell. (n) indicate the number of NCCs analyzed per condition. Error bars are s.e.m. (C) Scatter plots for the comparison of the number of PS-NH2 and PS-COOH NPs incorporated per cells after 20 h of exposure to NPs at each indicated concentration. PS-COOH data are shown in Fig 1B. One dot per cell. Error bars are s.e.m. (D) Proportion of NCCs having incorporated 0-5 PS-NH2 NPs (white bars), 6-50 PS-NH2 NPs (grey bars) or more than 50 PS-NH2 NPs (black bars) at 2h and 20 h, calculated from the data in B. Scale bars in the panels and insert in A are 25 and 50 μm, respectively. Data are from two independent experiments. Images in A are representatives from these experiments (n explants = 3). Data in B were compared between the different concentrations at each time point and between the two type points at each concentration and analyzed using non-parametric Kruskal-Wallis with uncorrected Dunn’s test, data in C were compared between PS-NH2 and PS-COOH NPs at each concentration and analyzed using non-parametric Kruskal-Wallis with uncorrected Dunn’s test, with only the significantly different P values <0.05 indicated; comparisons not indicated are not significant.

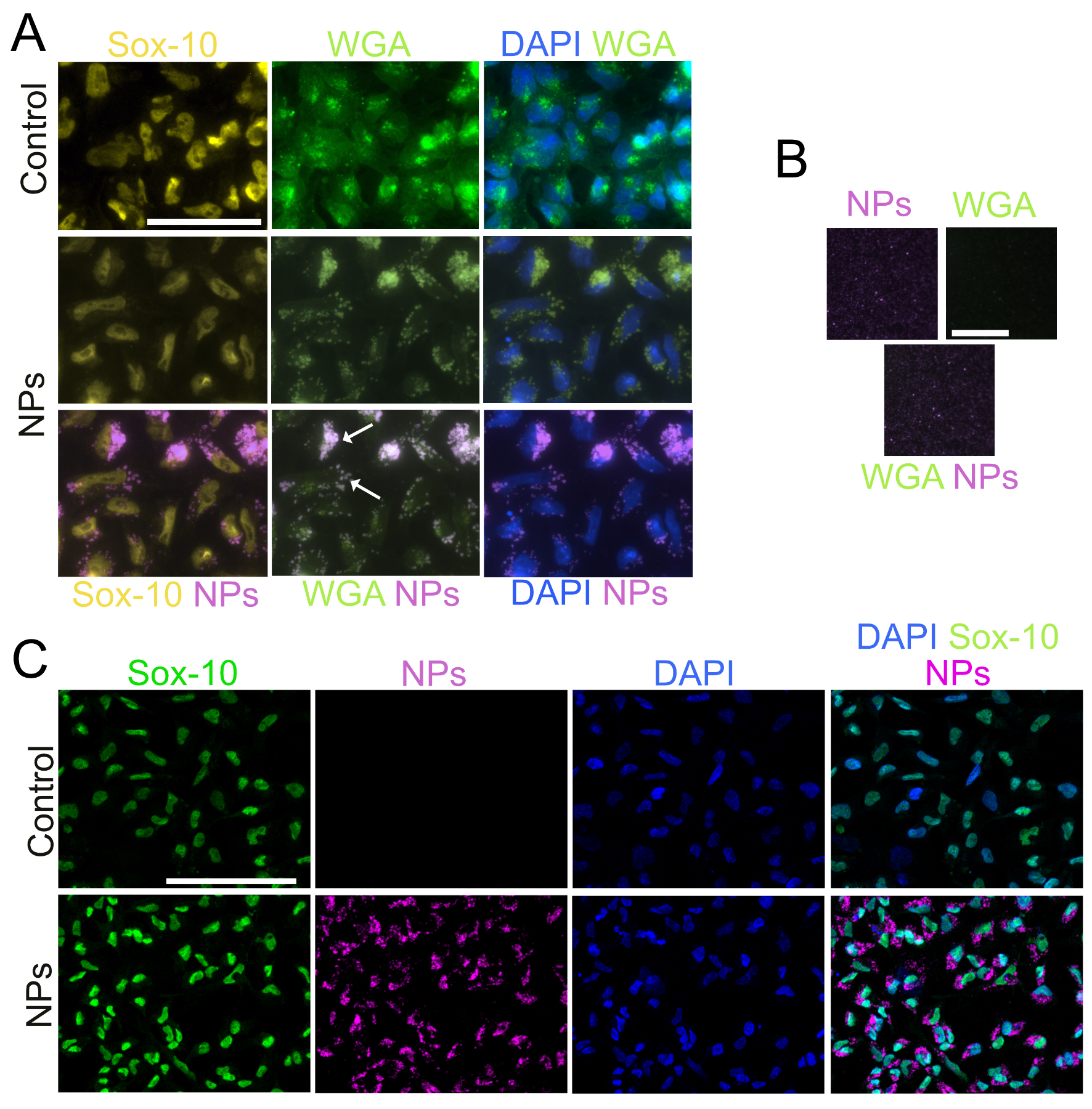

**S3 Fig, related to Fig 1. NCCs incorporated and accumulated 200nm NPs.** (A) Images showing the staining for wheat germ agglutinin (WGA, membranes, green), Sox-10 (NCC marker, nuclei, yellow), and with DAPI (nuclei, blue) in NCCs after 20 h of primary cultures in control condition (without NPs, top panels) and in the presence of NPs (middle and bottom panels) at 1X concentration, and merged images. Arrows point on NPs (magenta) incorporated in WGA positive structures. Scale bars, 50 μm. (B) Pictures taken showing that NPs (magenta) in cell-free zone of the field are not stained for WGA (green). Scale bars, 25 μm. (C) Images showing the staining for Sox-10 (NCC marker, nuclei, green), NPs (magenta), with DAPI (nuclei, blue), and merged images. Primary cultures were performed for 24 h in medium containing PS-COOH NPs at 10X concentration, then medium was rinsed and replaced by control medium for 24 h. Scale bars, 100 μm.

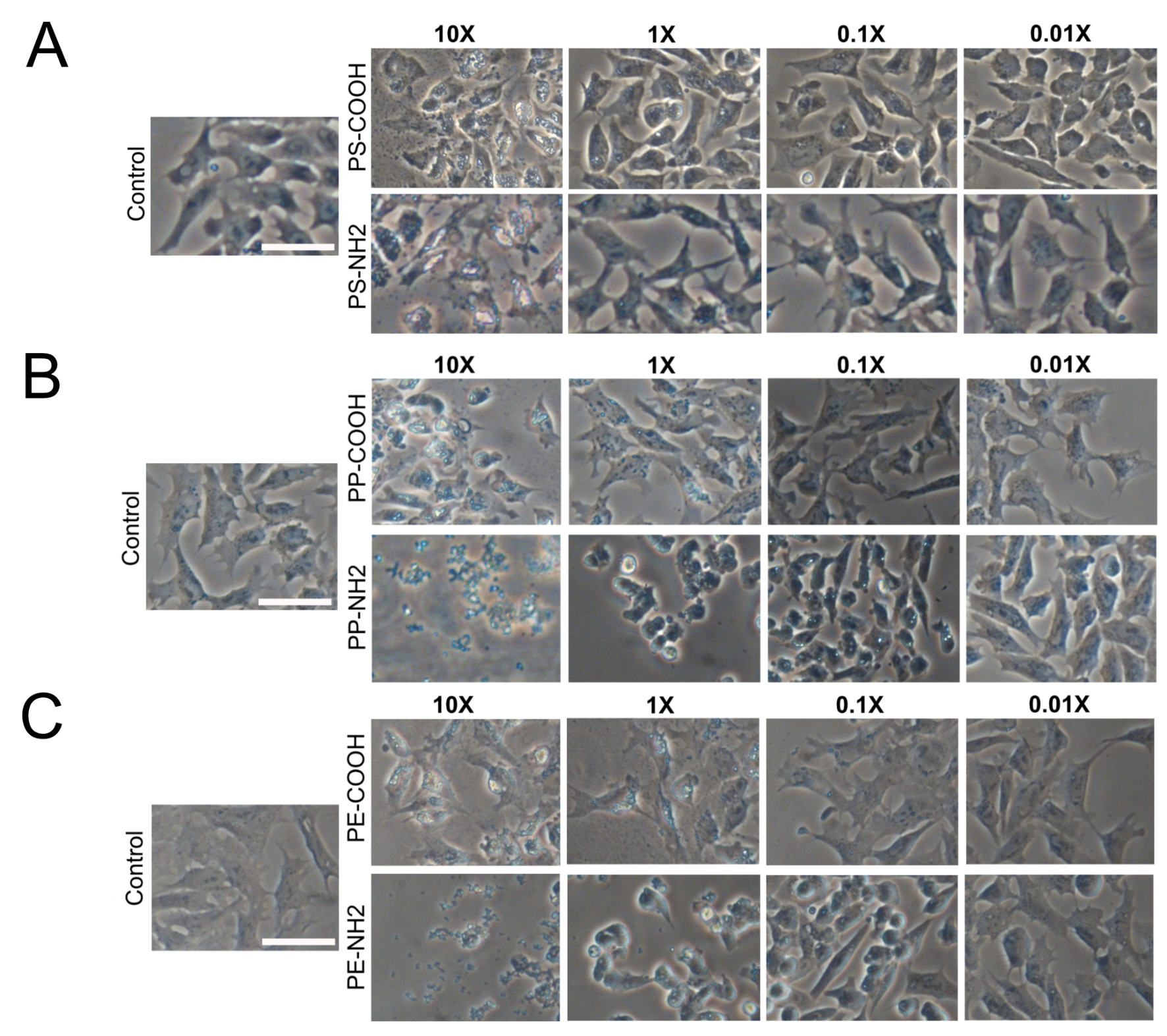

**S4 Fig, related to Figs 2 and 3. Effect of 200nm NPs on NCC morphology on primary cultures.** Phase contrast images at higher magnification of NCCs at 24 h primary cultures exposed to COOH- and NH2- forms of PS (A), PP (B), and PE (C) NPs at 0.01X, 0.1X, 1X and 10X concentrations, or in control condition (without NPs). Scale bar, 50 μm.

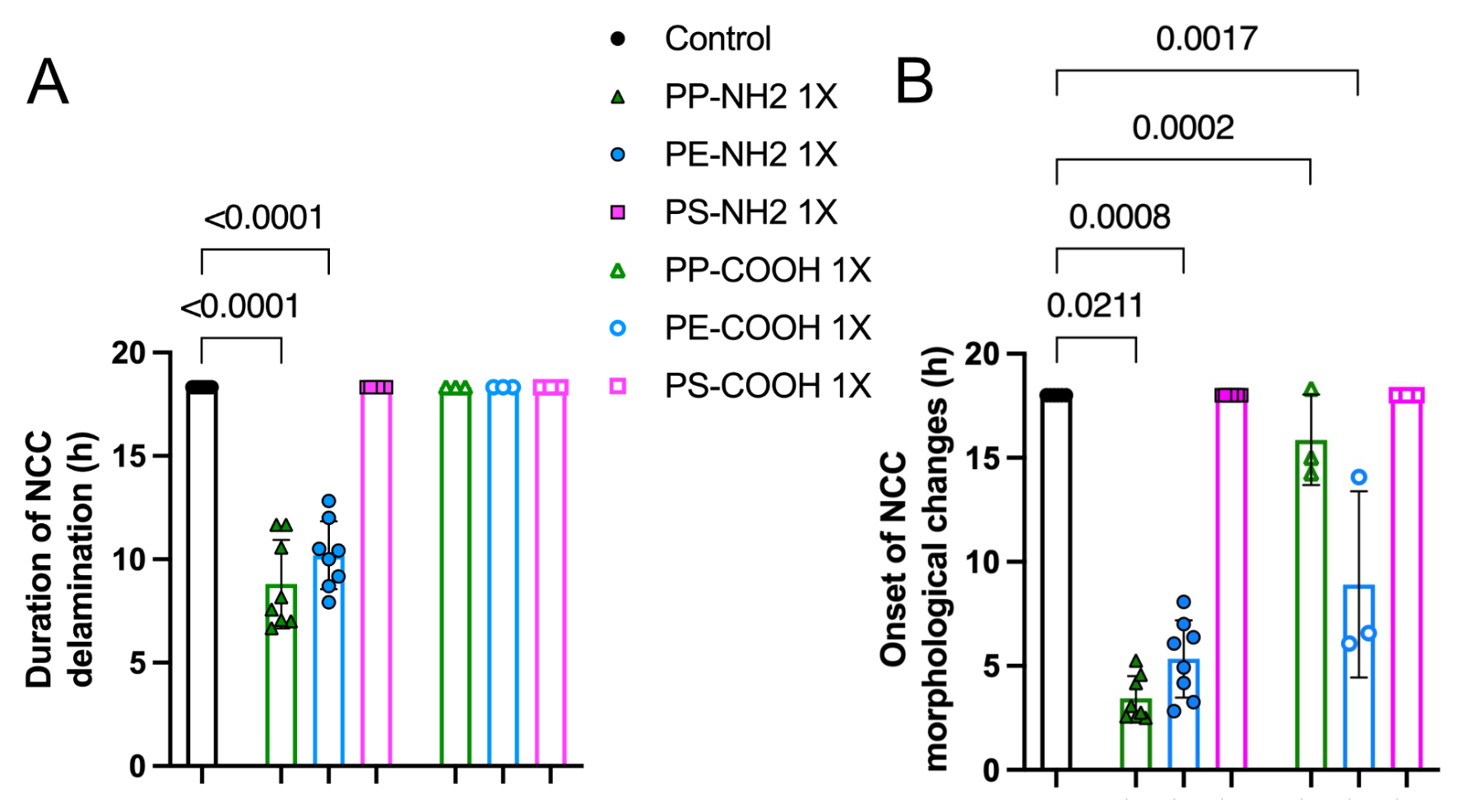

**S5 Fig, related to Figs 5 and 6. Effect of 200nm-NPs on NCC production time and the onset of morphological changes.** (A) Scatter plot of the NCC production period from neural tube in primary culture with mean ± S.D., measured from 18 hour-video recordings for cultures in the presence of NPs and in controls. When a neural tube was still producing NCCs at the end of the recording the value was set as 18h. Each dot represents one explant, with n explants = 8, 8, and 6, for PP-, PE- and PS-NH2 at 1X, respectively; n = 3 for each of NPs-COOH at 1X, and 8 for controls (pooled from the two independent experiments). (B) Scatter plot of the onset of NCC morphological changes in the outgrowth during migration compared that observed at t0 of video recording (starting 60 to 90 min after NPs were added to the cultures) with mean ± S.D., and in same conditions as in A. When no significant change in NCC morphology on cultures was observed during the 18-h recording the value was set as 18. Each dot represents one outgrowth, with n as in A. Data in A and B are obtained from the same recordings and are from two independent experiments. Data were analyzed using non-parametric Kruskal-Wallis with uncorrected Dunn’s test relative to the control, with only the significantly different P values <0.05 indicated; comparisons not indicated are not significant.

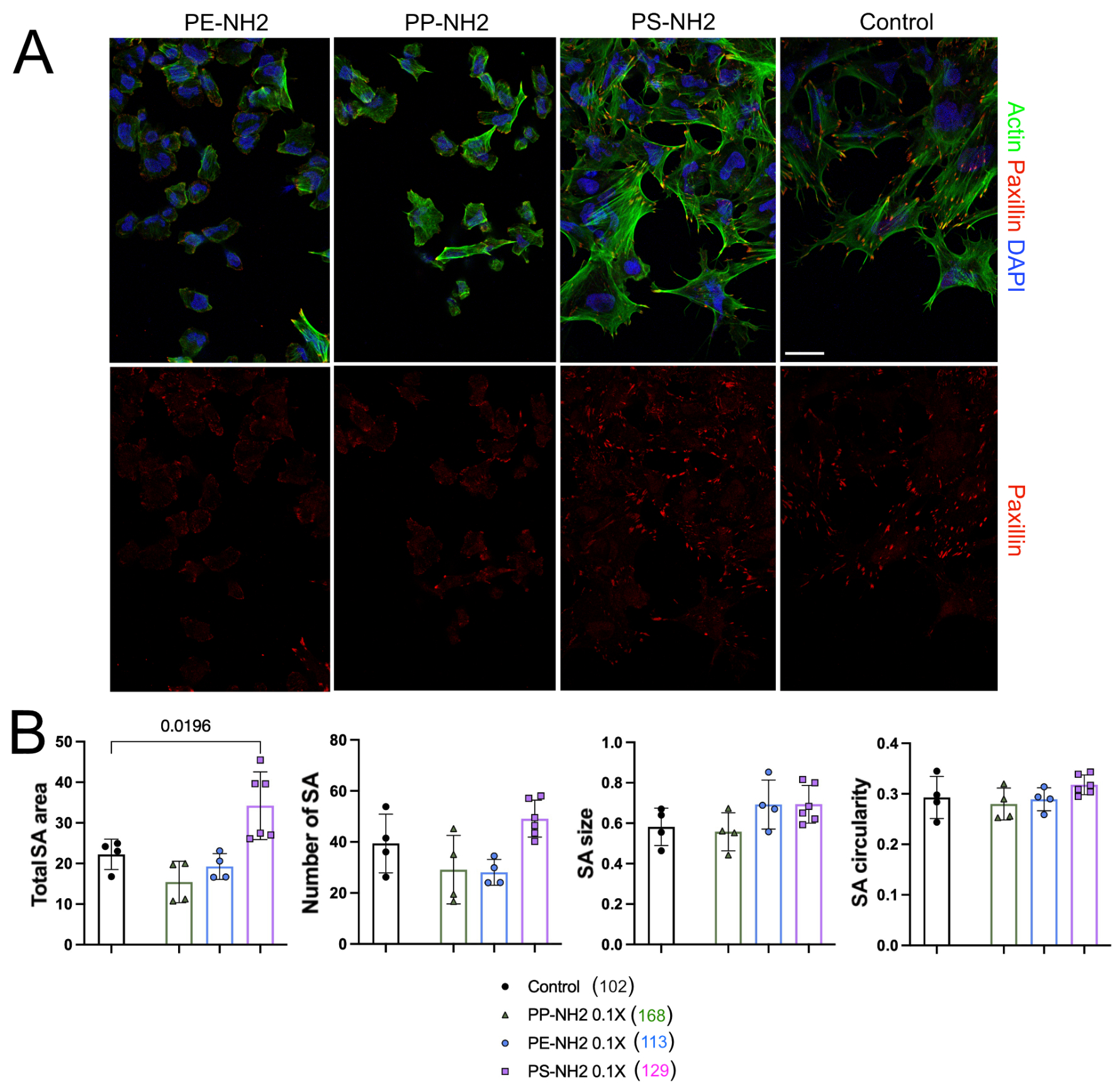

**S6 Fig, related to Fig 5. Effect of amine-functionalized 200nm NPs on NCC substrate adhesion.** (A) Immunostaining for Paxillin (red), phalloidin (green) and DAPI (blue, for nuclei) on NCCs cultured for 24 h in the presence of NPs-NH2 at 1X concentrations, and in control condition (without NPs). Scale bar, 20 μm. Images in A are representatives from the experiments quantified in Fig 5. (B) From left to right: Scatter plot of total substrate adhesion (SA) area per NCC, SA number, size and circularity, respectively, with mean ± S.D measured at 20 h of cultures in the presence of NPs-NH2 at 0.1X and in controls. Each dot represents the quantification from one image (n = 4, 4, and 6, for PP-, PE- and PS-NH2, respectively, and 4 for controls) randomly taken from the 2 different primary cultures. These conditions with NPs-NH2 0.1X were performed in the same experiments performed with NPs-COOH 1X and shared the controls shown in the Fig 6. (n) represent the total number of NCCs analyzed. Data were from two independent experiments. Images in A are representatives of these experiments. Data were analyzed by one-way ANOVA followed by Fisher's least significant difference (LSD) test for comparisons of each condition with the control, with only the significantly different P values <0.05 indicated; comparisons not indicated are not significant.

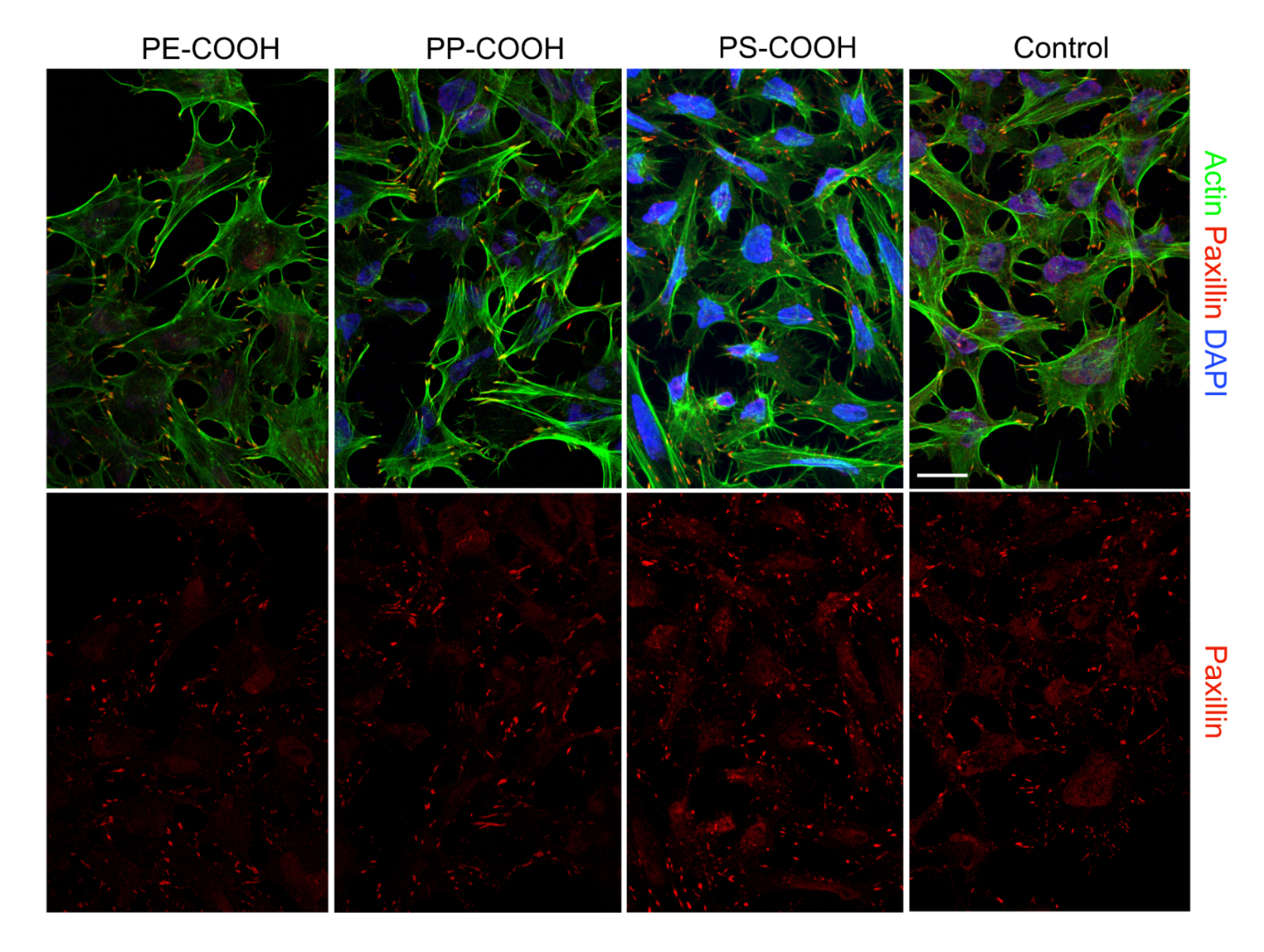

**S7 Fig, related to Fig 6. Effect of carboxylate-functionalized 200nm NPs on NCC substrate adhesion.** Immunostaining for Paxillin (red), phalloidin (green) and DAPI (blue, for nuclei) on NCCs cultured for 24 h in the presence of NPs-COOH at 1X concentrations, and in control condition. Scale bar, 20 μm. Images are representatives from the experiments quantified in Fig 6.
